# Fabrication of a Polymeric Inhibitor of Proximal Metabolic Enzymes in Hypoxia for Synergistic Inhibition of Cancer Cell Proliferation, Survival and Migration

**DOI:** 10.1101/2022.10.17.512469

**Authors:** Yuki Koba, Masahiko Nakamoto, Michiya Matsusaki

**Affiliations:** Division of Applied Chemistry, Graduate School of Engineering, Osaka University, Suita, Osaka 565-0871, Japan

**Keywords:** Polymer inhibitor, Enzyme inhibition, Cancer therapy, Multivalent interaction

## Abstract

Since conventional molecular targeted drugs often result in side effect, the development of novel molecular targeted drugs with both high efficacy and selectivity are desired. Simultaneous inhibition of metabolically and spatiotemporally related proteins/enzymes is a promising strategy for improving therapeutic interventions in cancer treatment. Herein, we report a poly-*α*-L-glutamate-based polymer inhibitor that simultaneously targets proximal transmembrane enzymes under hypoxia, namely carbonic anhydrase IX (CAIX) and zinc-dependent metalloproteinases. A polymer incorporating two types of inhibitors more effectively inhibited the proliferation and migration of human breast cancer cells than a combination of two polymers functionalized exclusively with either inhibitor. Synergistic inhibition of cancer cells would occur owing to the hetero-multivalent interactions of the polymer with proximate enzymes on the cancer cell membrane. Our results highlight the potential of polymer-based cancer therapeutics.

For Table of Contents only

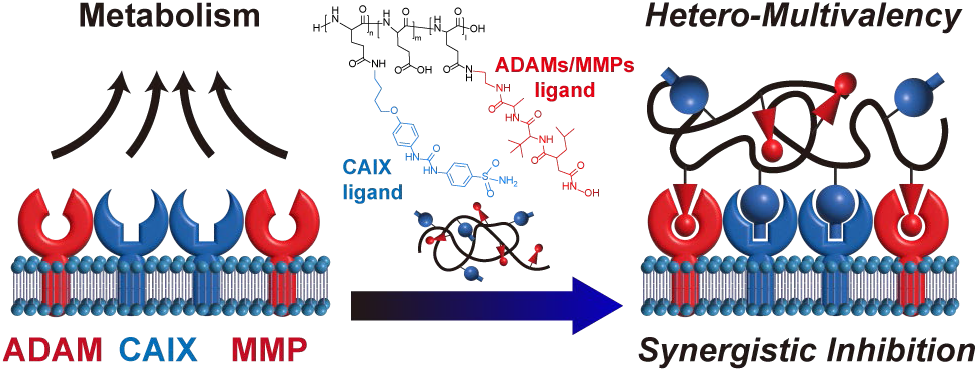

## INTRODUCTION

Molecular-targeted drugs that selectively inhibit target proteins have been developed for interventions in cancer therapy. However, drug discovery often results in disappointing returns owing to insufficient efficacy/selectivity which causes non-specific cytotoxicity and inhibition of isoforms and/or target proteins expressed on healthy cells.^1-4^ In addition, the depletion of therapeutic targets has also stagnated the recent drug discovery. Because the characteristic cancer cell metabolism including survival and proliferation is dependent on a limited number of molecular mechanisms, simultaneous inhibition of these vital mechanisms by a combination of drugs has been an attractive strategy for improving therapeutic intervention.^5-9^

Therapeutic efficacy can also be improved by the simultaneous inhibition of spatiotemporally co-localized proteins using the hetero-multivalent interactions.^10-12^ Lee *et al*. developed bispecific antibodies which showed effective antitumor properties by targeting the heterodimer consisting of epidermal growth factor receptor (EGFR) and hepatocyte growth factor receptor (Met).^13^ Polymers/nanoparticles, which display multiple ligands and recognize target biomacromolecules and/or cells by multivalent interactions,^14,15^ have been developed as abiotic antidotes,^16-19^ drugs,^20-23^ and carriers for drug delivery systems (DDS).^24,25^ Drug-free macromolecular therapeutic systems have recently been reported to induce apoptosis of human B lymphoblastoid cells by amplifying cross-linking of internalizing receptor CD20^26^ and to induce membrane disruption.^27^ DDS carriers with a ligand for the target proteins on a cancer cell have been widely studied for the effective delivery of anticancer drugs into the cells.^28^ Transferrin-conjugated nanoparticles loaded with camptothecin resulted in greater cellular uptake of drugs, sustaining intracellular drug retention, and enhancing the antitumor effect of the encapsulated agents.^29^ Moreover, the introduction of dual ligands in DDS carriers has improved their selective delivery and internalization efficiency.^30,31^ Hyaluronic acid-based nanogels targeting EGFR and cell-surface glycoprotein CD44 effectively delivered granzyme B with proliferative inhibition activity against breast cancer.^30^ The spatial control of ligands on DDS carriers has also been reported.^32,33^ A doxorubicin-loaded liposome-based nanocarrier, which was modified with targeting peptides on its fluidic surface, resulted in high specificity and cytotoxicity to target cells.^33^

However, to best our knowledge, there is no reported that polymer ligands that directly and synergistically inhibit cancer cell functions such as survival, proliferation, and migration via hetero-multivalent interactions with proximal enzymes on the cell membrane. We hypothesized that synergistic inhibition of cancer cells could be achieved by introducing two independent inhibitors for metabolically related and spatiotemporally proximal transmembrane enzymes in a polymer (Figure 1). To validate this hypothesis, carbonic anhydrase IX (CAIX) and related zinc-dependent metalloproteinases, including matrix metalloproteinases (MMPs) and a disintegrin and metalloproteinases (ADAMs), were selected as target enzymes. CAIX is a hypoxia-induced transmembrane enzyme that neutralizes intracellular pH acidified by glycolysis to promote tumor cell survival. CAIX inhibition has thus been regarded as a promising strategy for selective intervention in hypoxia-targeting cancer therapy.^34-36^ In addition, the combination of CAIX inhibition with anticancer drugs^37, 38^ and/or photosensitizers^39, 40^ has been exploited to enhance their anticancer efficacy. Recent interactome studies have revealed that zinc-dependent metalloproteinases such as MMP14 and ADAM17 are metabolically and spatiotemporally closely related to CAIX.^41-43^ MMP14, which promotes cancer cell metastasis and invasion by degrading the extracellular matrix, is activated by forming hetero-enzyme complexes with CAIX,^43^ while ADAM17 acts as a sheddase of CAIX for cellular signaling.^44,45^ Using a biocompatible poly (L-glutamic acid) as a scaffold, we report a polymer ligand functionalized with inhibitors for CAIX and MMPs/ADAMs which synergistically inhibits the proliferation, survival and migration of the highly aggressive human breast cancer cell line MDA-MB-231. This study provides a strategy to engineer the polymer inhibitor for cellular functions as well as the targeting ligand for DDS.

**Figure 1.**
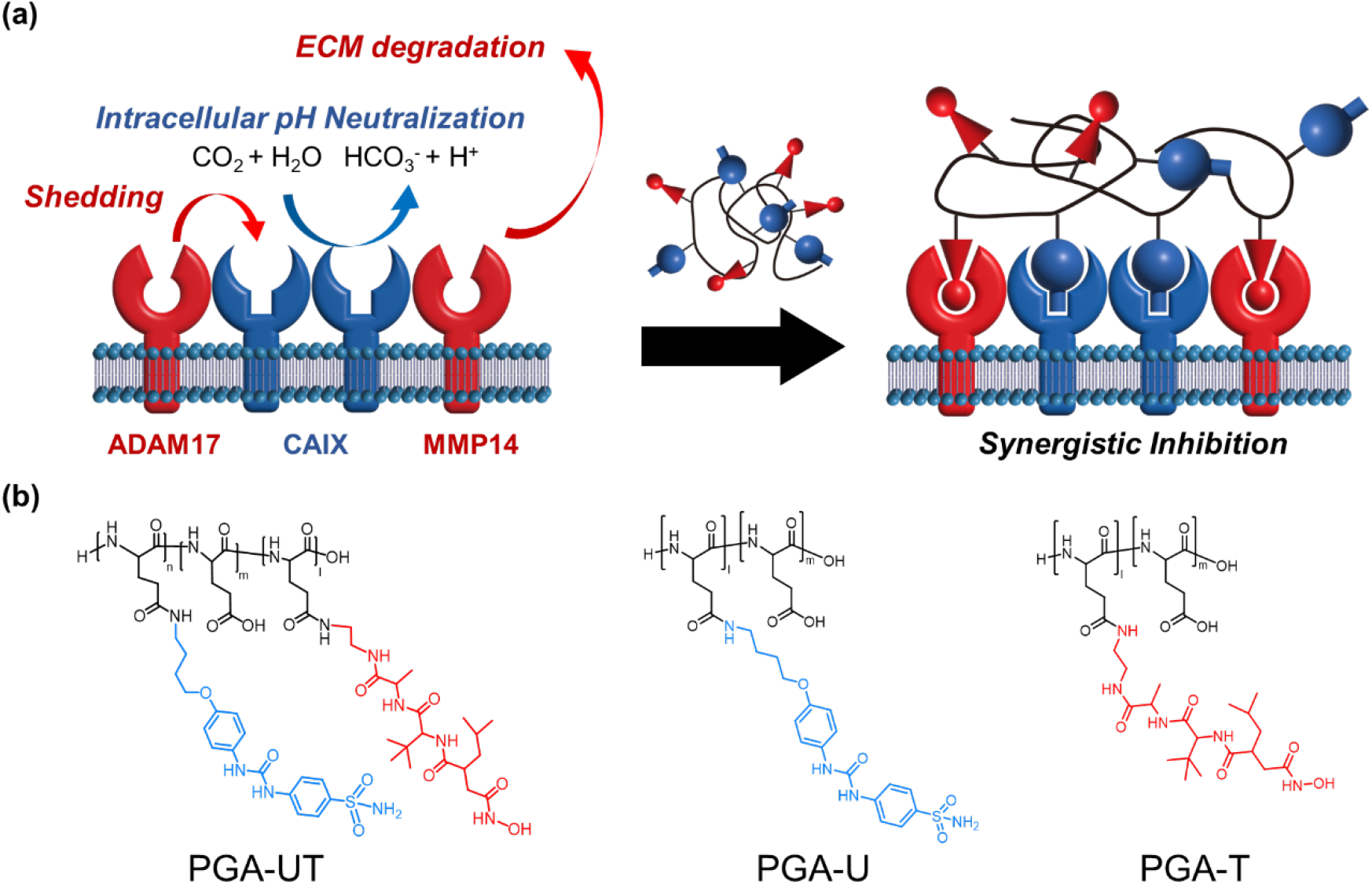
(a) Schematic illustration of CAIX- and MMPs/ADAMs-targeted poly (glutamic acid) (PGA) for synergistic inhibition of MDA-MB-231 human breast cancer cell proliferation, survival, and migration. (b) The chemical structure of PGA derivatives.

## EXPERIMENTAL SECTION

### Synthesis of PGA-UT

100 mM carbonate-bicarbonate buffer was prepared by mixing sodium carbonate (0.25 mmol) and sodium hydrogen carbonate in MilliQ (5 mL). Poly (L-glutamic acid) (417 nmol) was dissolved in 10 mM carbonate-bicarbonate buffer and sonicated for 10 min. The concentration of PGA derivatives in reaction mixture was 3 mg mL^−1^. Then, 500 mM 4-(4,6-dimethoxy-1,3,5-triazin-2-yl)-4-methylmorpholinium chloride (DMT-MM) aqueous solution was dropped into the solution and stirred for 10 min. (4-(3-(4-(4–amino butoxy)phenyl)ureido) benzene sulfonamide (U-104) aqueous solution was added into the reaction mixture and stirred for 24 h at room temperature. The reaction mixtures were dialyzed on a dialysis membrane in the following order: against 10 mM HCl for 1 day, against 10 mM NaOH for 1 day, and against MilliQ for 1 day, respectively, followed by lyophilized to obtain PGA-U. Yield = 100%. The grafting degree (G.D.) of PGA-U was calculated from UV-Vis absorbance at 270 nm relative to the calibration curve of U-104 (Figure S2). Then, 100 mM carbonate-bicarbonate buffer was prepared by mixing sodium carbonate (0.25 mmol) and sodium hydrogen carbonate in MilliQ (5 mL). PGA-U (41.7 nmol) was dissolved in 10 mM carbonate-bicarbonate buffer and sonicated 10 min. The concentration of PGA derivatives in eaction mixture was 3 mg mL^−1^. Then, 500 mM DMT-MM aqueous solution was dropped into the solution and stirred for 10 min. 40 mM *N*-(2-(2-(hydroxyamino)-2-oxoethyl)-4-methyl-1-oxopentyl)-3-methyl-L-valyl-*N*-(2-aminoethyl)-L-alaninamide (TAPI-2) ethanol solution was added into the reaction mixture and stirred for 7 h at room temperature. The reaction mixtures were dialyzed on a dialysis membrane against MilliQ for 3 days, followed by lyophilized to obtain PGA-UT. Yield = 100%. G.D. of TAPI-2 was calculated from ^1^H NMR spectrum (Figure S4, Table S1). The synthetic methods of PGA-T are shown in supporting information.

### Cell culture

Human breast cancer cell line (MDA-MB-231) and normal human dermal fibroblasts (NHDF) were obtained from KAC Co., Ltd. (Kyoto, Japan, EC92020424-F0) and Lonza (Basel, Switzerland, CC-2904), respectively. MDA-MB-231 cells were cultured in DMEM supplemented with 10% FBS and 1% Antibiotics. For culture under hypoxic conditions, cells were incubated at 37 °C under an atmosphere of 1% O_2_, 5% CO_2_, 94% N_2_ in a humidified incubator. For culture under normoxic conditions, cells were grown in a humidified incubator at 37 °C under 5% CO_2_.

### Water-soluble tetrazolium salts assay

The water-soluble tetrazolium salt assay (WST assay) was performed to quantitative cell viability treated with inhibitors and PGA derivatives. MDA-MB-231 cells (5.0 × 10^3^ cell/well) were seeded onto 96-well plates using DMEM adjusted to pH 6.8 by 1 M HCl and incubated for 24 h under 1% O_2_ (hypoxia). After incubation, cells were treated with inhibitors and PGA derivatives and incubated for 72 h under hypoxic conditions. Then, plates were washed with D-PBS and added cell counting kit-8 solution diluted 10 times by DMEM. After 3 h, plates were diluted 2 times by D-PBS and centrifuged (4000 rpm, 5 min). 80 µL supernatant were collected to 96-well plates and measured the absorbance at 450 nm using a micro plate reader. The IC_50_ values of PGA derivatives were calculated using GraphPad Prism (Figure S8).

### Visualization of the bonding of PGA derivatives on MDA-MB-231

The bonding of PGA derivatives on MDA-MB-231 was visualized using fluorescence-labelled PGA-U19T15 (PGA-UT-ATTO) and fluorescence-labelled PGA (PGA-ATTO) by confocal microscopy. The synthetic methods of fluorescence-labelled PGA derivatives are shown in supporting information (Figure S7, Table S1). Cells (3 × 10^4^ cell/well) were grown in 96-well glass bottom plate using DMEM adjusted to pH 6.8 by 1 M HCl and incubated for 24 h in 1% O_2_ (hypoxia). After incubation, PGA-UT-ATTO and PGA-ATTO was added to a final concentration of 100 nM, and cells were incubated for 1 h at 37 °C. The medium was removed, and after washing (3 × 100 µL DMEM), Hoechst 33342 (20 μg mL^−1^) was used to stain cell nuclei for 10 min. Confocal images were obtained with a FV 3000 confocal microscope and taken using a 405 nm laser diode (0.6%), for excitation with emission collected from 430 to 470 nm (detector voltage 720 V) for Hoechst 33342 and a 488 nm argon laser (0.5%) for excitation with emission collected from 506 to 606 nm (detector voltage 510 V) for ATTO-633. The fluorescence intensity of images was analyzed by the Imaris Software 9.0. The settings which were used here are shown in supporting information.

### Competitive Binding Inhibition Assay

MDA-MB-231 (3.0 × 10^4^ cell/well) were seeded into 96-well plate with glass bottom using DMEM adjusted to pH 6.8 by 1 M HCl and incubated for 24 h in 1% O_2_ (hypoxia). After incubation, mixture solution of U-104 (1 mM) and TAPI-2 (1 mM) was added and cells were incubated for 1 h at 37 ℃. PGA-UT-ATTO solution (16.7 µM) was added to a final concentration of 10 or 100 nM. After 1 h, the medium was removed, and after washing (1 × 100 µL DMEM), Hoechst 33342 (20 μg mL^−1^) was used to stain cell nuclei for 10 min. After washing (1 × 100 µL DMEM), confocal images were obtained with a FV 3000 confocal microscope and taken using a 405 nm laser diode (0.5%), for excitation with emission collected from 430 to 470 nm (detector voltage 550 V) for Hoechst 33342 and a 633 nm argon laser (0.5%) for excitation with emission collected from 506 to 606 nm (detector voltage 590 V) for ATTO-633. The fluorescence intensity of images was analyzed by the Imaris Software 9.0. The settings which were used here are shown in supporting information.

### Apoptosis/Necrosis/Healthy cell detection

MDA-MB-231 (5.0 × 10^3^ cell/well) were seeded into 96-well plates using DMEM adjusted to pH 6.8 by 1 M HCl and incubated for 24 h in 1% O_2_ (hypoxia). After incubation, cells were treated with PGA-U19T15 and incubated for 72 h in hypoxia. Then, following official procedure, Apoptosis/Necrosis/Healthy cell detection assay was performed with FV 3000. Images were taken using a 405 nm laser diode (0.5%), for excitation with emission collected from 430 to 470 nm (detector voltage 650 V) for Hoechst 33342, a 488 nm laser (0.5%) for excitation with emission collected from 500 to 552 nm (detector voltage 500 V) for FITC and a 561 nm laser (0.5%) for excitation with emission collected from 570 to 670 (detector voltage 510 V) for Ethidium homodimer-III. All images were taken using the same settings. Cell number of live cells, apoptosis, and necrosis were manually counted from images.

### Wound healing assay

MDA-MB-231 (3.0 × 10^4^ cell/well) were seeded into 96-well plates using DMEM adjusted to pH 6.8 by 1 M HCl and incubated for 24 h in hypoxia. Cells were treated with 5 μg/ml Mitomycin C for 2 h to inhibit the proliferation during the course of the assay, then wells were scratched by 200 µL pipet tips. The wound area of the cells growing in hypoxia were imaged at the zero and the 24 h time points with a phase contrast microscopy. Cell migration was calculated following equation:

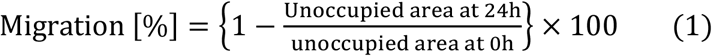

## RESULTS & DISCUSSION

### Synthesis of PGA derivatives

U-104^39,46^ and TAPI-2^47-49^ were used as an inhibitor for CAIX and MMPs/ADAMs, respectively. CAIX- and MMPs/ADAMs-targeted PGA derivatives were synthesized by functionalizing poly (L-glutamic acid) (PGA; Mw: 120 kDa) with U-104 and TAPI-2 via amide condensation using DMT-MM (Figure 1b, Scheme S2, Figure S2-5, and Table S1). Dynamic light scattering revealed that all PGA derivatives were soluble with a polymer size of approximately 10 nm under the experimental conditions and did not form large aggregates (Figure S6).

### Proliferative inhibitory effect of PGA derivatives

The proliferative inhibitory effect of PGA derivatives on the human breast cancer cell line (MDA-MB-231) under hypoxia (1% O_2_) and slightly acidic conditions (pH 6.8)^50^, in which CAIX and MMP/ADAM are overexpressed,^51^ was evaluated using a water-soluble tetrazolium salts assay (WST-8 assay).^52^ PGA without an inhibitor did not affect the cell proliferation, whereas PGA-U19 and PGA-T17 exclusively functionalized with either U-104 (G.D.: 19 mol%) or TAPI-2 (G.D.: 17 mol%) alone showed moderate proliferative inhibition (Figure 2a). Importantly, PGA-U19T15 conjugated with both U-104 and TAPI-2 (G.D. of U-104: 19 mol%; TAPI-2: 15 mol%) exhibited a higher proliferative inhibitory effect than that exerted by PGA-U19 and PGA-T17 alone. The results showed that the introduction of two inhibitors in PGA was responsible for the effective potency; the IC_50_ value of PGA-U19T15, PGA-U19, and PGA-T17 for proliferation were 1.33 µM, over 2.6, and over 2.9, respectively (Figure S8). The binding interface between the polymer and enzyme on the cell was further assessed by comparing the efficacy of PGA-U19 or PGA-T17 to that of small molecule inhibitor U-104 or TAPI-2 at the same inhibitor concentration (Figure 2b, c). The comparable efficacy of PGA-U19 or PGA-T17 with U-104 or TAPI-2 indicates that each polymer inhibits cell proliferation owing to monovalent binding of the inhibitor to the active site of the target enzyme, rather than multivalent interactions with a number of enzymes distributed on the cell membrane. Therefore, the efficacy of PGA-U19T15 with dual inhibitors would arise from its ability to inhibit proximal enzymes by hetero-multivalent interactions.

**Figure 2.**
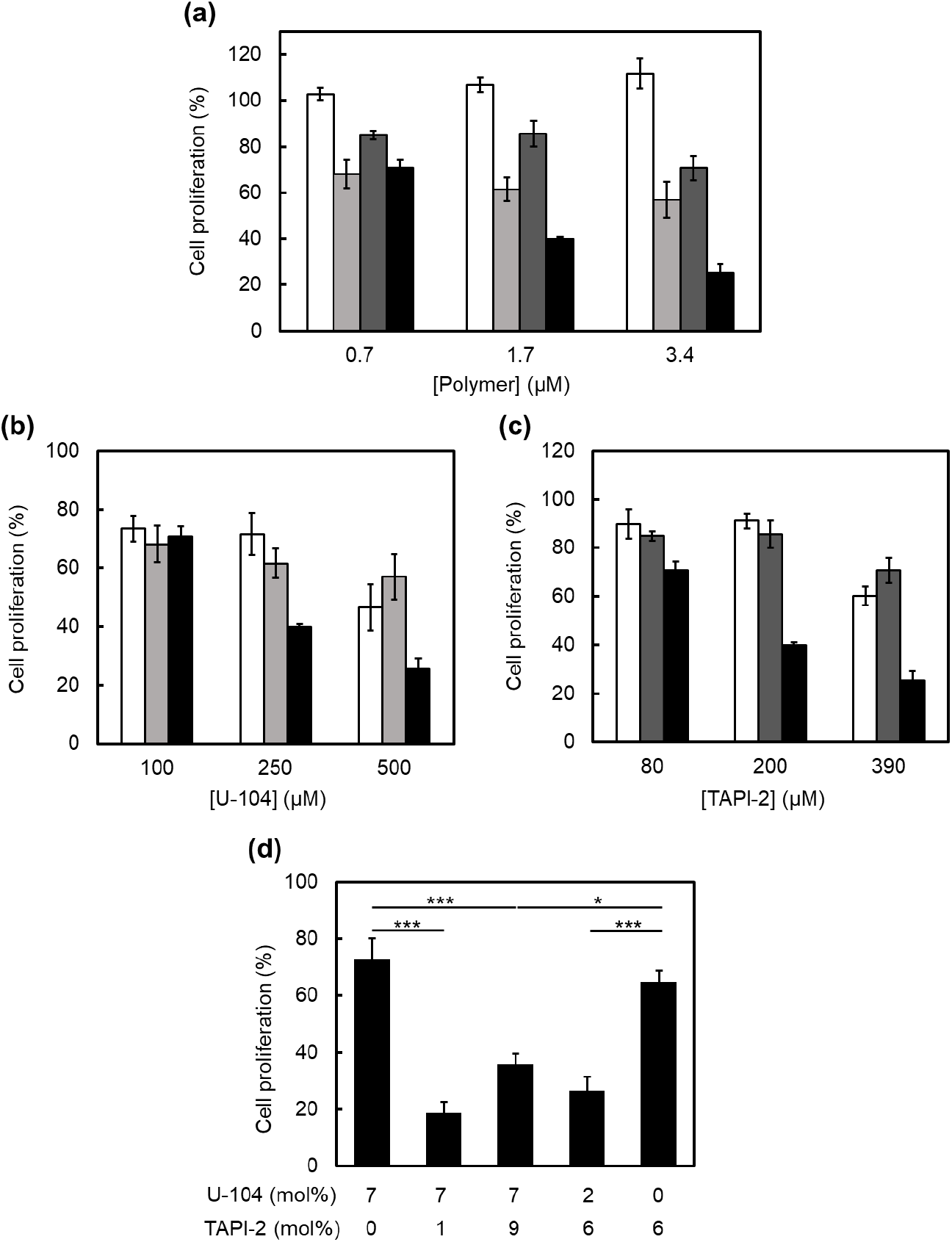
Cell proliferation of MDA-MB-231 treated with PGA derivatives, U-104 or TAPI-2 for 72 h under 1% O_2_ and pH 6.8 normalized to the proliferation of untreated cells (a) Cell proliferation after treatment with different polymer concentrations. White: PGA; light gray: PGA-U19; dark gray: PGA-T17; and black: PGA-U19T15. (b) Cell proliferation after treatment with different U-104 concentrations. White: U-104; light gray: PGA-U; and black: PGA-U19T15. (c) Cell proliferation after treatment with different TAPI-2 concentrations. White: TAPI-2; dark gray: PGA-T17; and black: PGA-U19T15. (d) Cell proliferation after treatment with PGA derivatives with different grafting degree of two inhibitors. Polymer concentration was 4.7 µM. Statistical analysis was performed using a two-tailed Student’s *t*-test from three independent experiments (n = 3). \**p* < 0.05 and \*\*\**p* < 0.005.

### Contribution of the amounts and proportion of two inhibitors into PGA towards the proliferative inhibitory effect

The efficacy of PGA-U7 with 7 mol% U-104 was significantly increased by incorporating 1 mol% TAPI-2, which was comparable with that of PGA-U19T15 (Figure 2d). Whereas, PGA-U7T9 with 7 mol% U-104 and 9 mol% TAPI-2 resulted in reduced efficacy. This result revealed that an increase in the multivalency of U-104 and TAPI-2 units in a polymer does not simply improve the efficacy because of the increased steric hindrance and/or decreased chain flexibility. In addition, when the total amount of inhibitors was fixed at 8 mol%, PGA-U7T1 with 7 mol% U-104 and 1 mol% TAPI-2 showed slightly lower cell proliferation than PGA-U2T6 with 2 mol% U-104 and 6 mol% TAPI-2. This result indicates that the effective inhibition might be achieved by optimizing the proportion of inhibitors in the polymer. Further investigations were performed using PGA-U19T15.

### Hypoxia selectivity of PGA-U19T15

Contrary to the significant efficacy of PGA-U19T15 in MDA-MB-231 cells under hypoxic conditions, PGA-UT showed marginal efficacy on MDA-MB-231 and normal human dermal fibroblasts (NHDF) under normoxic conditions, indicating the selective potency of PGA-U19T15 for the cancer cell under the hypoxic conditions in which CAIX and MMP/ADAM are overexpressed (Figure 3).

**Figure 3.**
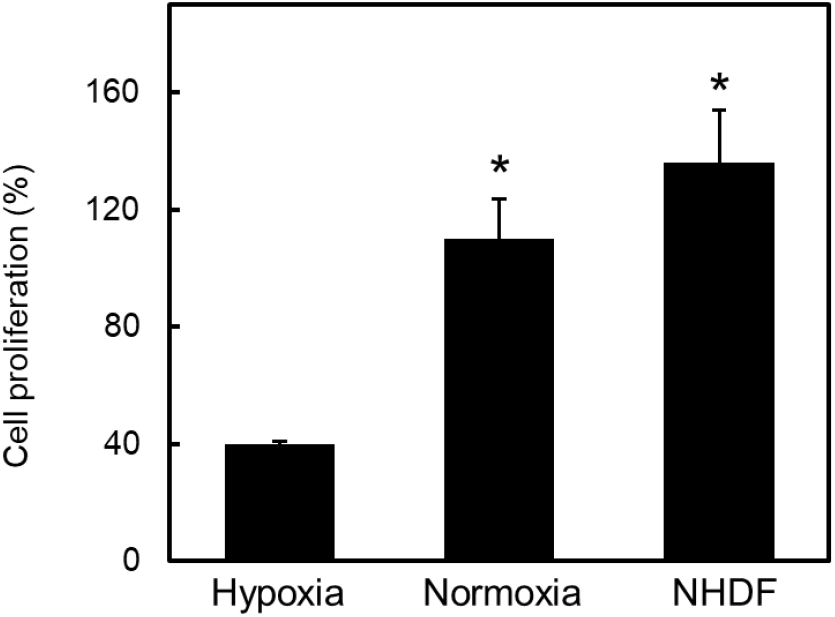
Cell proliferation of NHDF and MDA-MB-231 incubated at different oxygen concentration treated with PGA-U19T15 normalized to the proliferation of untreated cells. Polymer concentration was 1.7 µM. Statistical analysis was performed using a two-tailed Student’s *t*-test from three independent experiments (n = 3). \**p* < 0.05 relative to the hypoxic conditions.

### Targeting mechanism and a synergistic inhibitory effect of PGA-U19T15 to MDA-MB-231

The binding of polymers on MDA-MB-231 was visualized using fluorescence-labeled PGA-U19T15 (PGA-UT-ATTO) and PGA (PGA-ATTO) by introducing ATTO-633. Fluorescence imaging revealed the binding of PGA-UT-ATTO to cancer cells, whereas PGA-ATTO was not observed in the cells (Figure 4a). Confocal z slice images suggest the internalization of PGA-UT-ATTO in the cell (Figure S9c, d).^53,54^ Furthermore, fluorescence intensity from bound PGA-UT-ATTO on the cells decreased in the presence of a mixture of U-104 (1mM) and TAPI-2 (1 mM) (Figure 4b). Taken together, the results suggest that PGA-U19T15 selectively recognizes cancer cells by hetero-multivalent interaction with proximal enzymes comprising CAIX and MMPs/ADAMs overexpressed under hypoxic conditions, leading to effective suppression of cell proliferation. Furthermore, to validate the synergistic proliferative inhibitory effect of PGA-U19T15, the efficacy of PGA-U19T15 was compared to that of a mixture of small-molecule inhibitors (U-104+TAPI-2) and a mixture of PGA-U19 and PGA-T17 (PGA-U19+PGA-T17) at fixed concentrations of U-104 and TAPI-2 units. The efficacy of PGA-U19T15 was considerably higher than that of U-104+TAPI-2 and PGA-U19+PGA-T17 (Figure 4c). This result indicates that the introduction of two types of inhibitors in a polymer realizes synergistic inhibition of cancer cell proliferation owing to the hetero-multivalent interactions with proximal enzymes (Figure 4d). The insufficient efficacy of the mixture of PGA-U19 and PGA-T17 may be due to the uncooperative binding of each polymer to each target enzyme in addition to the steric hindrance of the binding of the two types of polymers to the proximal enzymes (Figure 4e).

**Figure 4.**
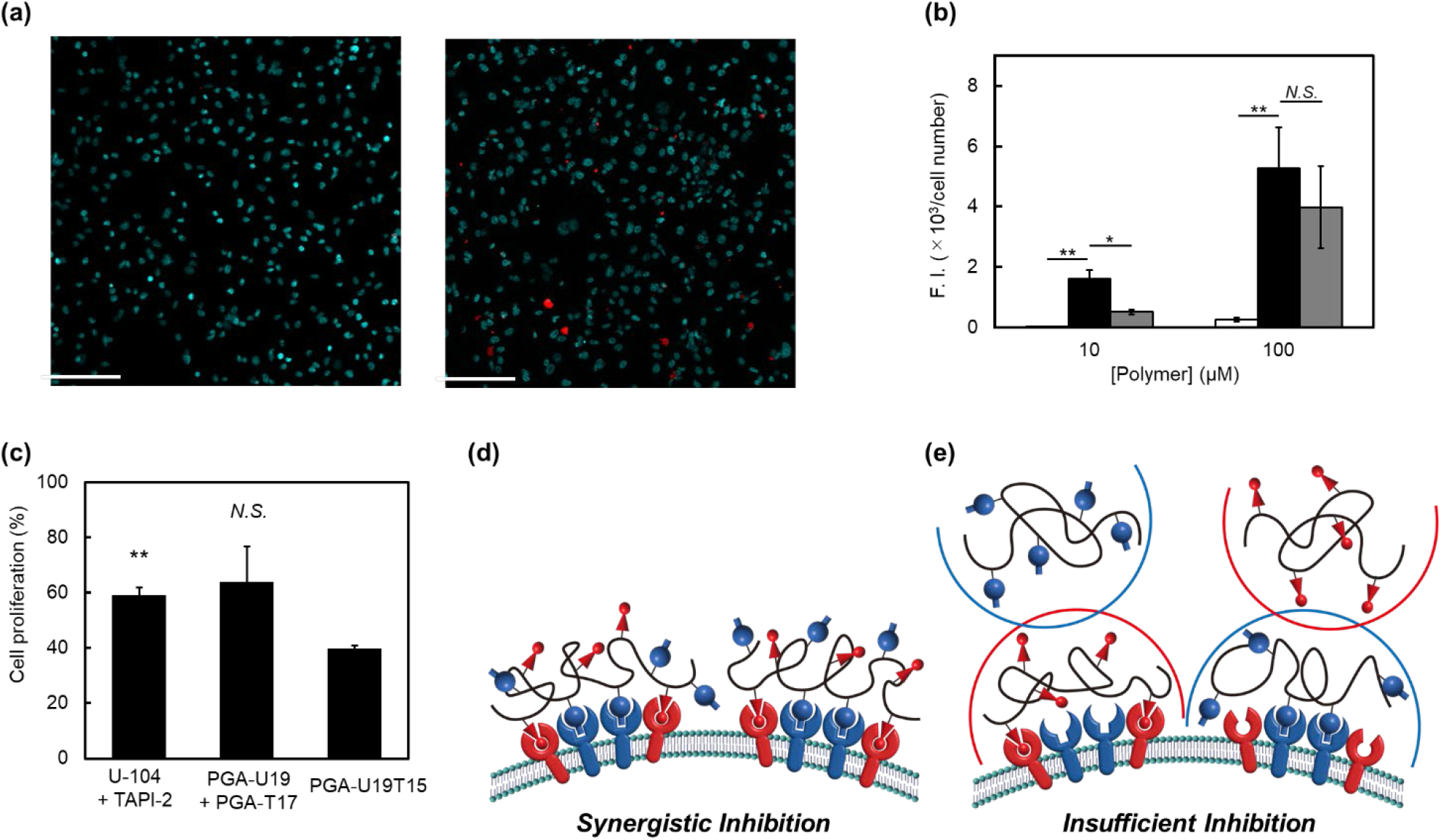
(a) Confocal images of MDA-MB-231 treated with labeled PGA (left) or PGA-U19T15 (right) for 1 h. Polymer concentration was 100 nM. Red: ATTO-633; blue: Hoechst 33342. Scale bar = 150 µm. (b) Fluorescence intensity from PGA derivatives normalized by cell number in images. White: PGA-ATTO; black: PGA-UT-ATTO in the absence of small-molecule inhibitors; gray: PGA-UT-ATTO in the presence of inhibitors. Statistical analysis was performed using a two-tailed Student’s *t*-test from three independent experiments (n = 3). \**p* < 0.05, \*\**p* < 0.01 relative to the PGA-U19T15 treated group. (c) Cell proliferation of MDA-MB-231 treated with PGA-U19T15, U-104+TAPI-2. and PGA-U19+PGA-T17 normalized to the proliferation of untreated cells. Concentrations of U-104 and TAPI-2 were fixed at 250 and 200 µM, respectively. Statistical analysis was performed using a two-tailed Student’s *t*-test from three independent experiments (n = 3). \*\**p* < 0.01 relative to the PGA-U19T15 treated group. (d, e) Schematic illustration of the inhibition of the proximal enzymes by (d) PGA-U19T15 and (e) PGA-U19+PGA-T17. Semi-circles indicate steric hindrance.

### Proliferative inhibition mechanism of PGA-U19T15 on MDA-MB-231

The mechanism of proliferative inhibition by PGA-U19T15 on MDA-MB-231 cells was investigated by staining apoptotic and necrotic cells (Figure 5a, b). PGA-U19T15 reduced the cell number by 33% compared to that of control cells without PGA-U19T15 treatment. Furthermore, the PGA-U19T15 treatment induced cell death: 4.9% of apoptotic and 8.1% of necrotic cells were observed; for the cells without treatment, 0.6% of apoptotic and 2.0% of necrotic cells were observed. Since it has been reported that CAIX inhibition induces the acidification of intracellular pH, resulting in cancer cell acidosis,^55^ PGA-U19T15 may induce the intracellular acidosis of cancer cells in the same manner.

**Figure 5.**
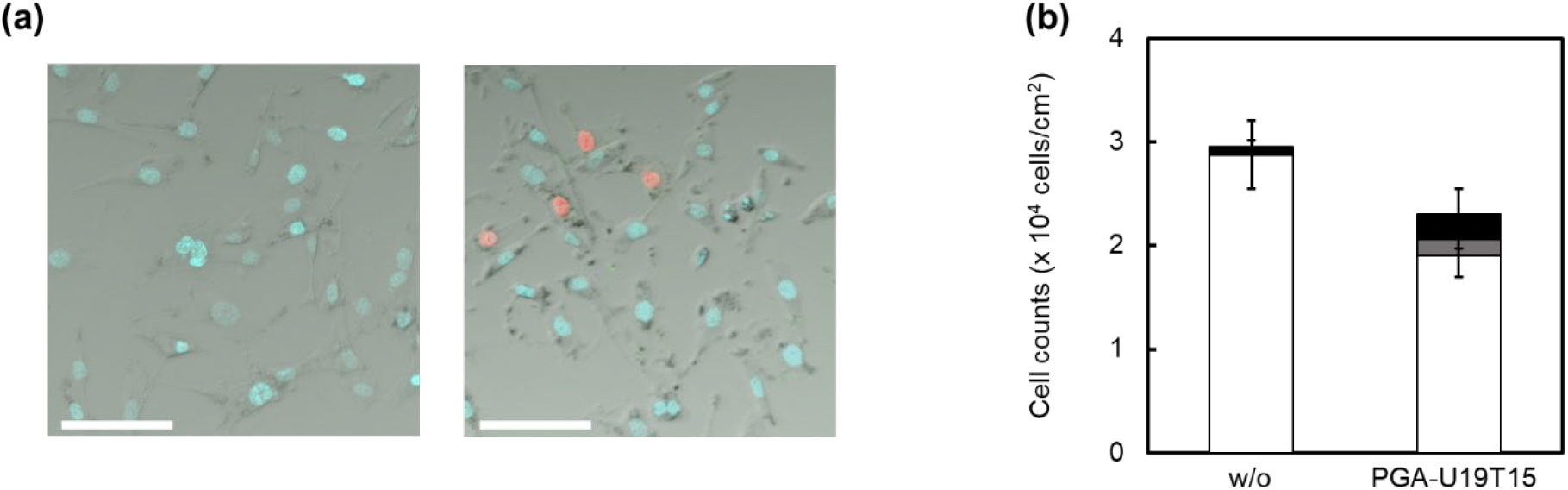
(a) Images of cells stained using apoptosis/necrosis/healthy cell detection kit. (Left: without PGA-U19T15 and right: treated with PGA-U19T15. Blue: live cells; green: apoptotic cells; and red: necrotic cells). Scale bar = 50 µm. (b) Quantification of the number of live cells, apoptosis, and necrosis using ImageJ software. The polymer concentration was 1.7 µM. White: live cells; gray: apoptotic cells; and black: necrotic cells.

### Migrative inhibitory effect of PGA-U19T15 on MDA-MB-231

We also performed a wound healing assay to demonstrate the inhibition of cell migration in MDA-MB-231 by PGA-U19T15. PGA derivatives inhibited MDA-MB-231 migration compared with that of the control cells (without PGA derivatives) (Figure 6a, b). Importantly, the migratory inhibition of MDA-MB-231 was significantly more effective with PGA-U19T15 than with PGA-U19 alone, PGA-T15 alone, and the mixture of PGA-U19 and PGA-T17 (Figure 6b). Furthermore, the mixture of inhibitors (U-104+TAPI-2) was more effective than PGA-U19T15, a result which may have arisen from the inhibition of other secreted MMPs including MMP2 and MMP9^56^ in the medium by small molecule TAPI-2.

**Figure 6.**
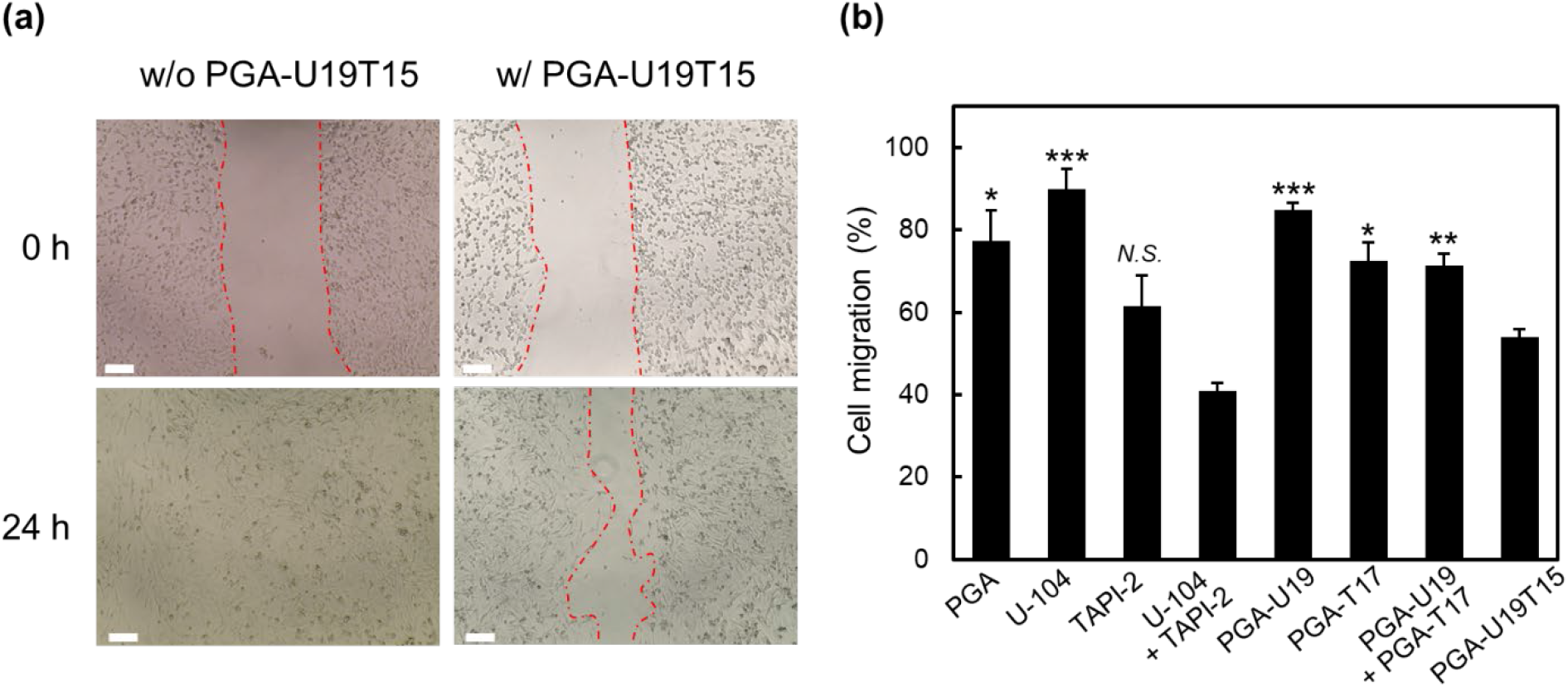
(a) Cell migration of MDA-MB-231 cells treated with inhibitors and PGA derivatives normalized to the proliferation of untreated cells. Polymer, U-104, and TAPI-2 concentrations were 1.7, 250, and 200 µM, respectively. Statistical analysis was performed using a two-tailed Student’s *t*-test from three independent experiments (n = 3). \**p* < 0.05, \*\**p* < 0.01, and \*\*\**p* < 0.005 relative to the PGA-U19T15 treated group. (b) Phase-contrast images of cell migration after 24 hours of incubation from scratching. Dashed lines demarcate wound boundaries. Left: cells treated without polymer and right: cells treated with PGA-U19T15. Top: t = 0 h and bottom: t = 24 h. Scale bar = 150 µm.

## CONCLUSIONS

In conclusion, we developed CAIX- and MMPs/ADAMs-targeted PGA derivatives functionalized with U-104 and TAPI-2, which significantly inhibit the proliferation, survival, and migration of MDA-MB-231 cells with the hypoxia selectivity. The results indicated that PGA-U19T15 synergistically inhibits the cellular function of MDA-MB-231 by targeting proximal enzymes owing to hetero-multivalent interactions. This study provides a strategy for developing polymer-based molecular-targeted drugs for synergistic inhibition of cancer cells, as well as the targeting ligand for a DDS system and/or the modulator that reduces the drug resistance of cancer cells.

## ASSOCIATED CONTENT

### Supporting Information

Additional data analysis includes ^1^H NMR spectra, UV-Vis spectra, DLS spectra and pictures.

## AUTHOR INFORMATION

### Corresponding Author

**Masahiko Nakamoto** - ^1^Division of Applied Chemistry, Graduate School of Engineering, Osaka University, Suita, Osaka 565-0871, Japan.

**Michiya Matsusaki** - ^1^Division of Applied Chemistry, Graduate School of Engineering, Osaka University, Suita, Osaka 565-0871, Japan.

## Author Contributions

The manuscript was written through contributions of all authors. All authors have given approval to the final version of the manuscript.

## ACKNOWLEDGMENT

This work was supported by Japan Society for the Promotion of Science (JSPS) Grant-in-Aid for Young Scientists (Start-up) (JP20K22499). This work was supported in part by JSPS Grant-in-Aid for Early-Career Scientists (JP22K18195), JSPS Scientific Research for Transformative Research Areas “Bottom-up Biotech” (JP22H05422) and Japan Science and Technology Agency (JST) COI-NEXT (JPMJPF2009).

## REFERENCE

(1) Lee, Y. T.; Tan, Y. J.; Oon, C. E. Molecular Targeted Therapy: Treating Cancer with Specificity. Eur. J. Pharmacol. 2018, 834, 188–196.

(2) Widakowich, C.; de Castro, G.; de Azambuja, E.; Dinh, P.; Awada, A. Review: Side Effects of Approved Molecular Targeted Therapies in Solid Cancers. Oncologist 2007, 12, 1443–1455.

(3) Vandenbroucke, R. E.; Libert, C. Is There New Hope for Therapeutic Matrix Metalloproteinase Inhibition? Nat. Rev. Drug Discov. 2014, 13, 904–927.

(4) Scannell, J. W.; Blanckley, A.; Boldon, H.; Warrington, B. Diagnosing the Decline in Pharmaceutical R&D Efficiency. Nat. Rev. Drug Discov. 2012, 11, 191–200.

(5) Narayan, R. S.; Molenaar, P.; Teng, J.; Cornelissen, F. M. G.; Roelofs, I.; Menezes, R.; Dik, R.; Lagerweij, T.; Broersma, Y.; Petersen, N.; Marin Soto, J. A.; Brands, E.; van Kuiken, P.; Lecca, M. C.; Lenos, K. J.; In ‘t Veld, S. G. J. G.; van Wieringen, W.; Lang, F. F.; Sulman, E.; Verhaak, R.; Baumert, B. G.; Stalpers, L. J. A.; Vermeulen, L.; Watts, C.; Bailey, D.; Slotman, B. J.; Versteeg, R.; Noske, D.; Sminia, P.; Tannous, B. A.; Wurdinger, T.; Koster, J.; Westerman, B. A. A Cancer Drug Atlas Enables Synergistic Targeting of Independent Drug Vulnerabilities. Nat. Commun. 2020, 11, 2935.

(6) Sawyers, C. L. Perspective: Combined Forces. Nature 2013, 498, S7–S7.

(7) Lehár, J.; Krueger, A. S.; Avery, W.; Heilbut, A. M.; Johansen, L. M.; Price, E. R.; Rickles, R. J.; Short, G. F.; Staunton, J. E.; Jin, X.; Lee, M. S.; Zimmermann, G. R.; Borisy, A. A. Synergistic Drug Combinations Tend to Improve Therapeutically Relevant Selectivity. Nat. Biotechnol. 2009, 27, 659–666.

(8) Hosaka, K.; Yang, Y.; Seki, T.; Du, Q.; Jing, X.; He, X.; Wu, J.; Zhang, Y.; Morikawa, H.; Nakamura, M.; Scherzer, M.; Sun, X.; Xu, Y.; Cheng, T.; Li, X.; Liu, X.; Li, Q.; Liu, Y.; Hong, A.; Chen, Y.; Cao, Y. Therapeutic Paradigm of Dual Targeting VEGF and PDGF for Effectively Treating FGF-2 off-Target Tumors. Nat. Commun. 2020, 11, 3704.

(9) Mokhtari, R. B.; Homayouni, T. S.; Baluch, N.; Morgatskaya, E.; Kumar, S.; Das, B.; Yeger, H. Combination Therapy in Combating Cancer. Oncotarget 2017, 8, 38022–38043.

(10) Becker, H. M.; Deitmer, J. W. Proton Transport in Cancer Cells: The Role of Carbonic Anhydrases. Int. J. Mol. Sci. 2021, 22, 3171.

(11) Becker, H. M. Carbonic Anhydrase IX and Acid Transport in Cancer. Br. J. Cancer 2020, 122, 157–167.

(12) Acunzo, M.; Romano, G.; Palmieri, D.; Laganá, A.; Garofalo, M.; Balatti, V.; Drusco, A.; Chiariello, M.; Nana-Sinkam, P.; Croce, C. M. Cross-Talk between MET and EGFR in Non-Small Cell Lung Cancer Involves MiR-27a and Sprouty2. Proc. Natl. Acad. Sci. 2013, 110, 8573.

(13) Lee, J. M.; Lee, S. H.; Hwang, J.-W.; Oh, S. J.; Kim, B.; Jung, S.; Shim, S.-H.; Lin, P. W.; Lee, S. B.; Cho, M.-Y.; Koh, Y. J.; Kim, S. Y.; Ahn, S.; Lee, J.; Kim, K.-M.; Cheong, K. H.; Choi, J.; Kim, K.-A. Novel Strategy for a Bispecific Antibody: Induction of Dual Target Internalization and Degradation. Oncogene 2016, 35, 4437–4446.

(14) Mammen, M.; Choi, S.-K.; Whitesides, G. M. Polyvalent Interactions in Biological Systems: Implications for Design and Use of Multivalent Ligands and Inhibitors. Angew. Chem. Int. Ed. Engl. 1998, 37, 2754–2794.

(15) Šácha, P.; Knedlík, T.; Schimer, J.; Tykvart, J.; Parolek, J.; Navrátil, V.; Dvořáková, P.; Sedlák, F.; Ulbrich, K.; Strohalm, J.; Majer, P.; ᘎubr, V.; Konvalinka, J. IBodies: Modular Synthetic Antibody Mimetics Based on Hydrophilic Polymers Decorated with Functional Moieties. Angew. Chem. Int. Ed. 2016, 55, 2356–2360.

(16) Hoshino, Y.; Koide, H.; Furuya, K.; Haberaecker, W. W.; Lee, S. H.; Kodama, T.; Kanazawa, H.; Oku, N.; Shea, K. J. The rational design of a synthetic polymer nanoparticle that neutralizes a toxic peptide in vivo. Proc. Natl. Acad. Sci. U. S. A. 2012, 109, 33−38.

(17) O’Brien, J.; Lee, S. H.; Onogi, S.; Shea, K. J. Engineering the Protein Corona of a Synthetic Polymer Nanoparticle for Broad-Spectrum Sequestration and Neutralization of Venomous Biomacromolecules. J. Am. Chem. Soc. 2016, 138, 16604−16607.

(18) Nakamoto, M.; Zhao, D.; Benice, O. R.; Lee, S. H.; Shea, K. J. Abiotic Mimic of Endogenous Tissue Inhibitors of Metalloproteinases: Engineering Synthetic Polymer Nanoparticles for Use as a Broad-Spectrum Metalloproteinase Inhibitor. J. Am. Chem. Soc. 2020, 142, 2338−2345.

(19) Pan, Y.; Yue, Y.; Hu, X.; Li, H.; Guo. D. A Supramolecular Antidote to Macromolecular Toxins Prepared through Coassembly of Macrocyclic Amphiphiles. Adv.Mater. 2021, 33, 2104310

(20) Koide, H.; Yoshimatsu, K.; Hoshino, Y.; Lee, S. H.; Okajima, A.; Ariizumi, S.; Narita, Y.; Yonamine, Y.; Weisman, A. C.; Nishimura, Y.; Oku, N.; Miura, Y.; Shea, K. J. A polymer nanoparticle with engineered affinity for a vascular endothelial growth factor (VEGF 165). Nat. Chem. 2017, 9, 715−722.

(21) Koide, H.; Okishima, A.; Hoshino, Y.; Kamon, Y.; Yoshimatsu, K.; Saito, K.; Yamauchi, I.; Ariizumi, S.; Zhou, Y.; Xiao, T.; Goda, K.; Oku, N.; Asail, T.; Shea, K. J. Synthetic hydrogel nanoparticles for sepsis therapy. Nat. Commun. 2021, 12, 5552.

(22) Yang, J.; Li, L.; Kopeček, J. Biorecognition: A key to drug-free macromolecular therapeutics. Biomaterials 2019, 190, 11–23.

(23) Raissi, A. J.; Scangarello, F. A.; Hulce, K. R.; Pontrello, J. K.; Paradis, S. Enhanced Potency of the Metalloprotease Inhibitor TAPI-2 by Multivalent Display. Bioorg. Med. Chem. Lett. 2014, 24, 2002–2007.

(24) Curk, T.; Dobnikar, J.; Frenkel, D. Optimal Multivalent Targeting of Membranes with Many Distinct Receptors. Proc. Natl. Acad. Sci. U. S. A. 2017, 114, 7210–7215.

(25) Rosenblum, D.; Joshi, N.; Tao, W.; Karp, J. M.; Peer, D. Progress and Challenges towards Targeted Delivery of Cancer Therapeutics. Nat. Commun. 2018, 9, 1410.

(26) Li, L.; Yang, J.; Wang, J.; Kopeček, J. Amplification of CD20 Cross-Linking in Rituximab-Resistant B-Lymphoma Cells Enhances Apoptosis Induction by Drug-Free Macromolecular Therapeutics. ACS Nano 2018, 12, 3658–3670.

(27) Nakatsuji, H.; Shioji, Y.; Hiraoka, N.; Okada, Y.; Kato, N.; Shibata, S.; Aoki, I.; Matsusaki, M. Cancer-Microenvironment Triggered Self-Assembling Therapy with Molecular Blocks. Mater. Horiz. 2021, 8, 1216–1221.

(28) Morales-Cruz, M.; Delgado, Y.; Castillo, B.; Figueroa, C. M.; Molina, A. M.; Torres, A.; Milián, M.; Griebenow, K. Smart Targeting To Improve Cancer Therapeutics. Drug Des. Dev. Ther. 2019, 13, 3753–3772.

(29) Hong, M.; Zhu, S.; Jiang, Y.; Tang, G.; Sun, C.; Fang, C.; Shi, B.; Pei, Y. Novel Anti-Tumor Strategy: PEG-Hydroxycamptothecin Conjugate Loaded Transferrin-PEG-Nanoparticles. J. Controlled Release 2010, 141, 22–29.

(30) Chen, J.; Ouyang, J.; Chen, Q.; Deng, C.; Meng, F.; Zhang, J.; Cheng, R.; Lan, Q.; Zhong, Z. EGFR and CD44 Dual-Targeted Multifunctional Hyaluronic Acid Nanogels Boost Protein Delivery to Ovarian and Breast Cancers In Vitro and In Vivo. ACS Appl. Mater. Interfaces 2017, 9, 24140–24147.

(31) Dong, Y.; Yu, T.; Ding, L.; Laurini, E.; Huang, Y.; Zhang, M.; Weng, Y.; Lin, S.; Chen, P.; Marson, D.; Jiang, Y.; Giorgio, S.; Pricl, S.; Liu, X.; Rocchi, P.; Peng, L. A Dual Targeting Dendrimer-Mediated SiRNA Delivery System for Effective Gene Silencing in Cancer Therapy. J. Am. Chem. Soc. 2018, 140, 16264–16274.

(32) Jiang, C.; Wang, X.; Teng, B.; Wang, Z.; Li, F.; Zhao, Y.; Guo, Y.; Zeng, Q. Peptide-Targeted High-Density Lipoprotein Nanoparticles for Combinatorial Treatment against Metastatic Breast Cancer. ACS Appl. Mater. Interfaces 2021, 13, 35248–35265.

(33) Ashley, C. E.; Carnes, E. C.; Phillips, G. K.; Padilla, D.; Durfee, P. N.; Brown, P. A.; Hanna, T. N.; Liu, J.; Phillips, B.; Carter, M. B.; Carroll, N. J.; Jiang, X.; Dunphy, D. R.; Willman, C. L.; Petsev, D. N.; Evans, D. G.; Parikh, A. N.; Chackerian, B.; Wharton, W.; Peabody, D. S.; Brinker, C. J. The Targeted Delivery of Multicomponent Cargos to Cancer Cells by Nanoporous Particle-Supported Lipid Bilayers. Nat. Mater. 2011, 10, 389–397.

(34) Alterio, V.; Hilvo, M.; Di Fiore, A.; Supuran, C. T.; Pan, P.; Parkkila, S.; Scaloni, A.; Pastorek, J.; Pastorekova, S.; Pedone, C.; Scozzafava, A.; Monti, S. M.; De Simone, G. Crystal Structure of the Catalytic Domain of the Tumor-Associated Human Carbonic Anhydrase IX. Proc. Natl. Acad. Sci. U. S. A. 2009, 106, 16233–16238.

(35) Pastorekova, S.; Gillies, R. J. The Role of Carbonic Anhydrase IX in Cancer Development: Links to Hypoxia, Acidosis, and Beyond. Cancer Metastasis Rev. 2019, 38, 65–77.

(36) Wilson, W. R.; Hay, M. P. Targeting Hypoxia in Cancer Therapy. Nat. Rev. Cancer 2011, 11, 393–410.

(37) Kuijk, S. J. A. van; Gieling, R. G.; Niemans, R.; Lieuwes, N. G.; Biemans, R.; Telfer, B. A.; Haenen, G. R. M. M.; Yaromina, A.; Lambin, P.; Dubois, L. J.; Williams, K. J. The Sulfamate Small Molecule CAIX Inhibitor S4 Modulates Doxorubicin Efficacy. PLoS One 2016, 11, e0161040.

(38) Cao, Q.; Zhou, D.-J.; Pan, Z.-Y.; Yang, G.-G.; Zhang, H.; Ji, L.-N.; Mao, Z.-W. CAIXplatins: Highly Potent Platinum (IV) Prodrugs Selective Against Carbonic Anhydrase IX for the Treatment of Hypoxic Tumors. Angew. Chem. 2020, 132, 18715–18721.

(39) Pospíšilová, K.; Knedlík, T.; Šácha, P.; Kostka, L.; Schimer, J.; Brynda, J.; Král, V.; Cígler, P.; Navrátil, V.; Etrych, T.; Šubr, V.; Kugler, M.; Fábry, M.; Řezáčová, P.; Konvalinka, J. Inhibitor–Polymer Conjugates as a Versatile Tool for Detection and Visualization of Cancer-Associated Carbonic Anhydrase Isoforms. ACS Omega 2019, 4, 6746–6756.

(40) Kim, J. H.; Verwilst, P.; Won, M.; Lee, J.; Sessler, J. L.; Han, J.; Kim, J. S. A Small Molecule Strategy for Targeting Cancer Stem Cells in Hypoxic Microenvironments and Preventing Tumorigenesis. J. Am. Chem. Soc. 2021, 143, 14115–14124.

(41) McDonald, P. C.; Swayampakula, M.; Dedhar, S. Coordinated Regulation of Metabolic Transporters and Migration/Invasion by Carbonic Anhydrase IX. Metabolites 2018, 8, E20.

(42) Zatovicova, M.; Sedlakova, O.; Svastova, E.; Ohradanova, A.; Ciampor, F.; Arribas, J.; Pastorek, J.; Pastorekova, S. Ectodomain Shedding of the Hypoxia-Induced Carbonic Anhydrase IX Is a Metalloprotease-Dependent Process Regulated by TACE/ADAM17. Br. J. Cancer 2005, 93, 1267–1276.

(43) Swayampakula, M.; McDonald, P. C.; Vallejo, M.; Coyaud, E.; Chafe, S. C.; Westerback, A.; Venkateswaran, G.; Shankar, J.; Gao, G.; Laurent, E. M. N.; Lou, Y.; Bennewith, K. L.; Supuran, C. T.; Nabi, I. R.; Raught, B.; Dedhar, S. The Interactome of Metabolic Enzyme Carbonic Anhydrase IX Reveals Novel Roles in Tumor Cell Migration and Invadopodia/MMP14-Mediated Invasion. Oncogene 2017, 36, 6244–6261.

(44) Vidlickova, I.; Dequiedt, F.; Jelenska, L.; Sedlakova, O.; Pastorek, M.; Stuchlik, S.; Pastorek, J.; Zatovicova, M.; Pastorekova, S. Apoptosis-Induced Ectodomain Shedding of Hypoxia-Regulated Carbonic Anhydrase IX from Tumor Cells: A Double-Edged Response to Chemotherapy. BMC Cancer 2016, 16, 239.

(45) Zunke, F.; Rose-John, S. The Shedding Protease ADAM17: Physiology and Pathophysiology. Biochim. Biophys. Acta, Mol. Cell Res. 2017, 1864, 2059–2070.

(46) McDonald, P. C.; Chia, S.; Bedard, P. L.; Chu, Q.; Lyle, M.; Tang, L.; Singh, M.; Zhang, Z.; Supuran, C. T.; Renouf, D. J.; Dedhar, S. A Phase 1 Study of SLC-0111, a Novel Inhibitor of Carbonic Anhydrase IX, in Patients With Advanced Solid Tumors. Am. J. Clin. Oncol. 2020, 43, 484–490.

(47) Kruse, M.-N.; Becker, C.; Lottaz, D.; Köhler, D.; Yiallouros, I.; Krell, H.-W.; Sterchi, E. E.; Stöcker, W. Human Meprin Alpha and Beta Homo-Oligomers: Cleavage of Basement Membrane Proteins and Sensitivity to Metalloprotease Inhibitors. Biochem. J. 2004, 378, 383–389.

(48) Wang, R.; Ye, X.; Bhattacharya, R.; Boulbes, D. R.; Fan, F.; Xia, L.; Ellis, L. M. A Disintegrin and Metalloproteinase Domain 17 Regulates Colorectal Cancer Stem Cells and Chemosensitivity Via Notch1 Signaling. Stem Cells Transl. Med. 2016, 5, 331–338.

(49) Hooper, N. M.; Karran, E. H.; Turner, A. J. Membrane Protein Secretases. Biochem. J. 1997, 321, 265–279.

(50) Warburg, O.; Wind, F.; Negelein, E. THE METABOLISM OF TUMORS IN THE BODY. J. Gen. Physiol. 1927, 8, 519–530.

(51) Meehan, J.; Ward, C.; Turnbull, A.; Bukowski-Wills, J.; Finch, A. J.; Jarman, E. J.; Xintaropoulou, C.; Martinez-Perez, C.; Gray, M.; Pearson, M.; Mullen, P.; Supuran, C. T.; Carta, F.; Harrison, D. J.; Kunkler, I. H.; Langdon, S. P. Inhibition of PH Regulation as a Therapeutic Strategy in Hypoxic Human Breast Cancer Cells. Oncotarget 2017, 8, 42857–42875.

(52) Peskin, A. V.; Winterbourn, C. C. A Microtiter Plate Assay for Superoxide Dismutase Using a Water-Soluble Tetrazolium Salt (WST-1). Clinica. Chimica. Acta 2000, 293, 157–166.

(53) Christianson, H. C.; Menard, J. A.; Chandran, V. I.; Bourseau-Guilmain, E.; Shevela, D.; Lidfeldt, J.; Månsson, A.-S.; Pastorekova, S.; Messinger, J.; Belting, M. Tumor Antigen Glycosaminoglycan Modification Regulates Antibody-Drug Conjugate Delivery and Cytotoxicity. Oncotarget 2017, 8, 66960–66974.

(54) Zatovicova, M.; Jelenska, L.; Hulikova, A.; Ditte, P.; Ditte, Z.; Csaderova, L.; Svastova, E.; Schmalix, W.; Boettger, V.; Bevan, P.; Pastorek, J.; Pastorekova, S. Monoclonal Antibody G250 Targeting CA ?: Binding Specificity, Internalization and Therapeutic Effects in a Non-Renal Cancer Model. Int. J. Oncol. 2014, 45, 2455–2467.

(55) Temiz, E.; Koyuncu, I.; Durgun, M.; Caglayan, M.; Gonel, A.; Güler, E. M.; Kocyigit, A.; Supuran, C. T. Inhibition of Carbonic Anhydrase IX Promotes Apoptosis through Intracellular PH Level Alterations in Cervical Cancer Cells. Int. J. Mol. Sci. 2021, 22, 6098.

(56) Hegedüs, L.; Cho, H.; Xie, X.; Eliceiri, G. L. Additional MDA-MB-231 Breast Cancer Cell Matrix Metalloproteinases Promote Invasiveness. J. Cell. Physiol. 2008, 216, 480–485.

