## Supporting Information for "Fabrication of a Polymeric Inhibitor of Proximal Metabolic Enzymes in Hypoxia for Synergistic Inhibition of Cancer Cell Proliferation, Survival and Migration"

Table of contents

1. Materials

2. Synthesis of U-104

3. Synthesis of PGA derivatives

4. WST assay

5. Visualization of the binding of PGA-UT to MDA-MB-231

### 1. Materials

#### Reagents

The following materials were obtained from commercial sources.

Tetrahydrofuran (207-17765), 5 mol/L sodium hydroxide solution (1310-73-2), potassium hydrogen sulfate (7646-93-7), magnesium sulfate (10034-99-8), toluene super dehydrated (108-88-3), acetonitrile super dehydrated (75-05-8), 5 mol/L hydrochloric acid (7647-01-0), 4 M Hydrogen chloride・ethyl acetate solution (083-10405), sodium carbonate (497-19-8), sodium hydrogen carbonate (144-55-8), *N*, *N*- dimethylformamide (68-12-2), phosphate buffer saline (D-PBS; D5652-10L), and trypsin (9002-07-7) were obtained from Wako, Japan. Diisopropyl azodicarboxylate (2446-83-5), dimethyl sulfoxide-D6 (2206-617-0), deuterium oxide (7789-20-0) and mitomycin C from *Streptomyces caespitosus* (50-07-7) were obtained from Sigma Aldrich, U.S.A. Chloroform-D1 0.03vol% TMS (865-49-6) was obtained from Merck, Germany. Methyl 4-hydroxy benzoate (99-76-3), triphenylphosphine (603-35-0), *N*, *N*-diisopropylethylamine (7087-68-5), diphenyl phosphoryl azide (26386-88-9) and sulfanilamide (63-74-1) were obtained from Tokyo Chemical Industry Co., Ltd., Japan. Sodium deuteroxide (D, 99.5%) 40% in $D_2O$ (14014-063) was obtained from Cambridge Isotope Laboratories, Inc., U.S.A. n-Hexane (110-54-3), ethyl acetate (EtOAc; 141-78-6), methanol (67-56-1) and ethylenediaminetetraacetic acid (EDTA; 10378-23-1) were obtained from KISHIDA CHEMICAL, Japan. Poly (L-glutamic acid) (PGA; 26247-79-0) was obtained from Alamanda

Polymers, U.S.A. 4- (*tert*-Butoxycarbonylamino)-1-butanol (75178-87-9) was obtained from Combi Blocks, U.S.A. DMT-MM・$nH_2O$ (2170798-10-2) was obtained from WATANABE CHEMICAL, Japan. TAPI-2 (187034-31-7) was obtained from Med Chem Express, U.S.A. Dulbecco's modified Eagle's medium (DMEM; 08458-45) and Cell Count Reagent sf (07553-44) were obtained from nacalai tesque, Japan. Fatal bovine serum (FBS; 10270), trypan blue (2309124), NHS-rhodamine (246256-50-8) and Hoechst 33342 (23491-52-3, H3570) were purchased from Thermo Fisher Scientific, U.S.A. Apoptosis/Necrosis/Healthy cell detection kit (PK-CA707-30018) was obtained from PromoCell GmbH, Germany. ATTO-633 amine (AD633-91) was obtained from ATTO-TECH GmbH, Germany. Atelocollagen acidic solution IPC-30 was obtained from KOKEN, Japan.

**Instrumentation**

$^1$H NMR spectra and ESI-MS spectra were measured using JNM-GSX 400 (JEOL, Japan) and JMS-T100LP (JEOL, Japan), respectively. UV-Vis and fluorescence spectra were recorded using NanoDrop™ 2000c (Thermo Fisher Scientific, U.S.A.) and FP-8500 spectrofluorometer (JASCO, Japan), respectively. Dynamic light scattering was performed using Zetasizer Nano ZS (Malvern, U.S.A). Polymers were purified by dialysis using Spectra/Por® 7 Dialysis Membrane (MWCO: 8 kDa, Repligen, U.S.A.). Polymers were sonicated using sonicator US-105 (SND, Japan). Solution pH was measured by LAQUA twin pH-22B (Horiba, Japan). Phase-contrast images and confocal images were taken using EVOS XL core (Thermo Fisher Scientific, U.S.A.) and FV3000 (Olympus, Japan), respectively. Confocal images were analyzed using Imaris Cell Imaging Software 9.0 (Bitplane, UK). The $IC_{50}$ value was calculated by GraphPad Prism (GraphPad Software, U.S.A.).

### 2. Synthesis of CAIX inhibitor U-104

**Scheme S1.** Synthesis of CAIX inhibitor U-104

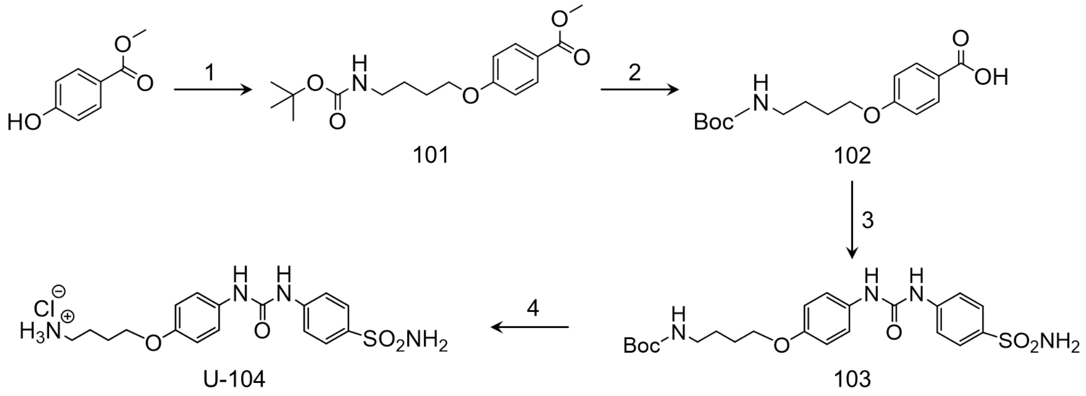


We synthesized U-104, CAIX inhibitor, on the basis of the reported procedure (Scheme S1).[39]

#### 2-1. Synthesis of compound 101

Methyl 4-hydroxybenzoate (35.3 mmol) and triphenylphosphine (52.9 mmol) were dissolved in THF dehydrated (210 mL). Then, 4- (*tert*-Butoxycarbonylamino)-1-butanol (52.9 mmol) dissolved in THF dehydrated (10 mL) was added into the solution. Diisopropyl azodicarboxylate (52.9 mmol) was dropped to the reaction solution. The reaction was stirred overnight under a nitrogen atmosphere at room temperature. The reaction mixture was evaporated to remove solvent and the crude product was centrifuged after addition of hexane/ethyl acetate (3 : 1) mixed solution (10000 rpm, 10 min). Supernatant was purified by column chromatography (hexane: ethyl acetate = 3: 1, Rf = 0.25-0.40) and evaporated to yield a white powder (10.1g). Yield = 88%. $^{1}$H NMR (400 MHz, $CDCl_3$): δ 7.98 (d, J = 8.4 Hz, 2H), 6.90 (d, J = 8.4 Hz, 2H), 4.60 (s, 1H), 4.03 (t, J = 6.2 Hz, 2H), 3.88 (s, 3H), 3.20 (m, 2H), 1.87-1.80 (m, 2H), 1.71-1.64 (m, 2H), 1.45 (s, 9H). MS (ESI+): calculated for $C_{17}H_{25}O_5N$ $[MNa]^+$; 346.17. Found, 346.2.

**2-2. Synthesis of compound 102**

Compound 1 (17.4 mmol) was dissolved in methanol (104.6 mL) and 5 M NaOH (104.6 mL) was added. The reaction mixture was left stirring for 3 h at 50 °C. The reaction was traced by thin-layer chromatography (TLC) analysis (hexane: ethyl acetate = 3 : 1, Rf of compound 2 = 0). The reaction mixture was evaporated to remove methanol, then, diluted with ethyl acetate (250 mL). The aqueous phase was acidified with 10% $KHSO_4$ and extracted twice with ethyl acetate (100 mL). We checked the existence of compound 2 in the water phase with TLC (hexane: ethyl acetate = 3: 1) but there was no spot showing compound 2. Then, the oil phase was evaporated and dried in a vacuum to yield a white powder (4.12 g). Yield = 72%. $^{1}$H NMR (400 MHz, $CDCl_3$): δ 8.04 (d, J = 9.1 Hz, 2H), 6.92 (d, J = 9.1 Hz, 2H), 4.61 (s, 1H), 4.05 (t, J = 6.3 Hz, 2H), 3.21 (m, 2H), 1.89-1.82 (m, 2H), 1.71 (m, 2H), 1.45 (s, 9H) MS (ESI-): calculated for $C_{16}H_{22}O_5N$ $[M]^-$; 308.16. Found, 308.2.

**2-3. Synthesis of compound 103**

Compound 102 (2.3 mmol) was dissolved in toluene dehydrates (65 mL), and *N*, *N*-diisopropylethylamine (4.65 mmol) was added under a nitrogen atmosphere at 90 °C. Under the same condition, diphenyl phosphoryl azide (2.56 mmol) was added to the reaction mixture in one portion and the mixture was stirred for 5 h. The reaction was traced by thin-layer chromatography (TLC) analysis (hexane: ethyl acetate = 2:5, Rf of the product = 0.35). The reaction mixture was then evaporated and dissolved in acetonitrile hydrated (30 mL), then, the mixture was heated to 60 °C. Sulfanilamide (3.49 mmol, Rf = 0.36) was added in one portion, then, the reaction mixture was stirred overnight under a nitrogen atmosphere. The reaction mixture was evaporated and dried in a vacuum for 5 h. The crude product was dissolved in ethyl acetate (60 mL), washed 10 times with 30 mM HCl (50 mL). Then the organic layer was evaporated to obtain white precipitation. In this washing process, we confirmed that compound 3 was in the oil layer with TLC (hexane: ethyl acetate = 2: 5, Rf = 0.25). The precipitates were collected by filtration using a Kiriyama funnel and dried in vacuum to obtain a white powder (2.59 g). Yield = 42%. $^{1}$H NMR (400 MHz, DMSO-$d_6$): δ 8.98 (s, 1H), 8.59 (s, 1H), 7.71 (d, J = 8.7 Hz, 2H), 7.59 (d, J = 9.2 Hz, 2H), 7.34 (d, J = 9.2 Hz, 2H), 7.19 (s, 2H), 6.86 (d, J = 8.7 Hz, 3H), 3.91 (t, J = 6.4 Hz, 2H), 2.96 (q, J = 6.4 Hz, 2H), 1.70-1.63 (m, 2H), 1.51 (q, J = 7.3 Hz, 2H), 1.37 (s, 9H).

**2-4. Synthesis of U-104**

Compound 3 (2.26 mmol) was dissolved in THF (5 mL) under a nitrogen atmosphere, then, sonicated for 10 min. The mixture was added 4 M HCl・EtOAc (6.5 mL) and stirred for 90 min at room temperature. The reaction mixture was evaporated and dried in a vacuum to yield an off-white powder. Yield = 96%.[1]H NMR (400 MHz, DMSO- $d_6$): 9.53 (s, 1H), 9.11 (s, 1H), 7.84 (s,3H), 7.71 (d, J = 8.6 Hz, 2H), 7.59 (d, J = 8.6 Hz, 2H), 7.37-7.32 (m, 2H), 7.19 (s, 2H), 6.89-6.83 (m, 2H), 3.94 (q, J = 5.9 Hz, 2H), 2.84 (d, J = 4.5 Hz, 2H), 1.75-1.69 (m, 4H).

**3. Synthesis of PGA derivatives**

**Scheme S2.** Synthesis of PGA derivatives.

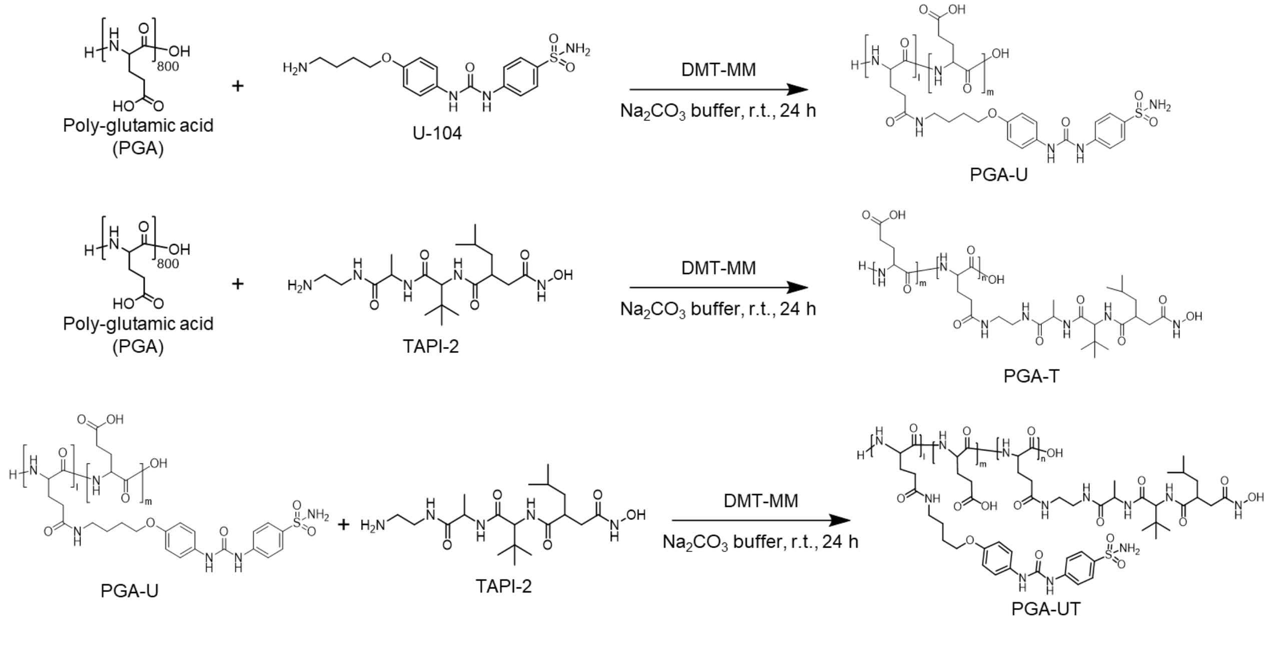


**3-1. Evaluation of reactivity of the terminal amine group of poly (L-glutamic acid)**

It was necessary for the reaction of PGA with inhibitors to confirm the reactivity of the terminal amine group of PGA. Synthesis of PGA-Rhodamine was performed to evaluate the reactivity of them. 100 mM carbonate-bicarbonate buffer was prepared by mixing sodium carbonate (0.25 mmol) and sodium hydrogen carbonate in ultrapure water (MilliQ) (5 mL). Poly (glutamic acid) (20.8 nmol) was dissolved in 10 mM carbonate-bicarbonate buffer (50 μL) and sonicated 10 min. 18.9 mM NHS-rhodamine DMF solution (246 nmol) was dropped into the solution and the reaction mixture was stirred for 2 h on ice bath. The reaction mixtures were dialyzed on a dialysis membrane against MilliQ for 1 days, followed by lyophilized to obtain PGA-Rho (2.3 mg). Grafting degree of rhodamine was calculated to be 2% from fluorescence intensity at 579 nm. The results indicated that the reactivity of erminal amine group of PGA would be neglectable during amide condensation of PGA and inhibitors.

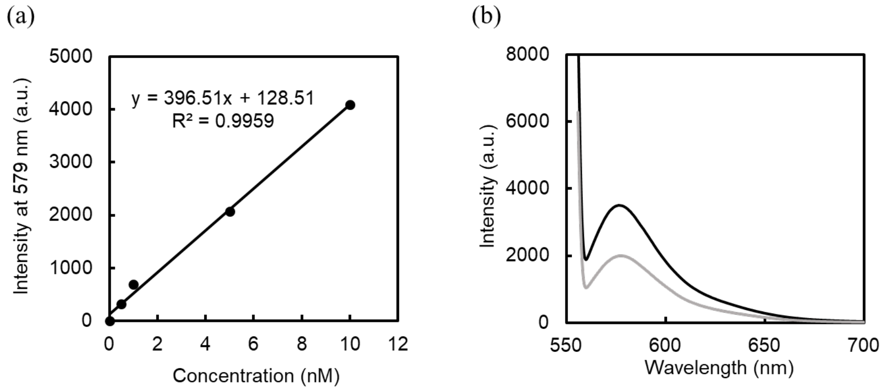


**Figure S1.** Fluorescence measurement in MilliQ. Excitation: 552 nm and Fluorescent: 579 nm. (a) The calibration curve of NHS-rhodamine. (b) Fluorescence spectra of PGA-Rho (black: 10 eq. 415 μM and gray: 2.5 eq. 913 nM).

#### 3-2. Synthesis of PGA-U

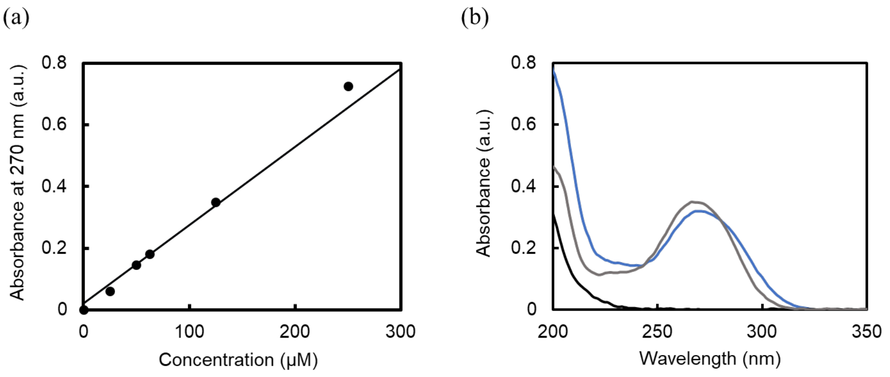


**Figure S2.** UV-Vis measurement in MilliQ. (a) The calibration curve of U-104. (b) UV-Vis spectra of PGA (black, 0.15 mg $mL^{-1}$), U-104 (gray, 125 μM), and PGA-U (blue, 1 mg $mL^{-1}$).

#### 3-3. Synthesis of PGA-T

100 mM carbonate-bicarbonate buffer was prepared by mixing sodium carbonate (0.25 mmol) and sodium hydrogen carbonate in MilliQ (5 mL). Poly (glutamic acid) (417 nmol) was dissolved in 10 mM carbonate-bicarbonate buffer and sonicated 10 min. The concentration of PGA derivatives in reaction mixture was 3 mg $mL^{-1}$. Then, 500 mM DMT-MM aqueous solution was dropped into the solution and stirred for 10 min. 40 mM TAPI-2 ethanol solution was added into the reaction mixture and stirred for 24 h at room temperature. The reaction mixtures were dialyzed on a dialysis membrane against MilliQ for 3 days, followed by lyophilized to obtain PGA-T. Yield = 100%. Grafting degree of TAPI-2 was calculated from $^{1}$H NMR spectrum.

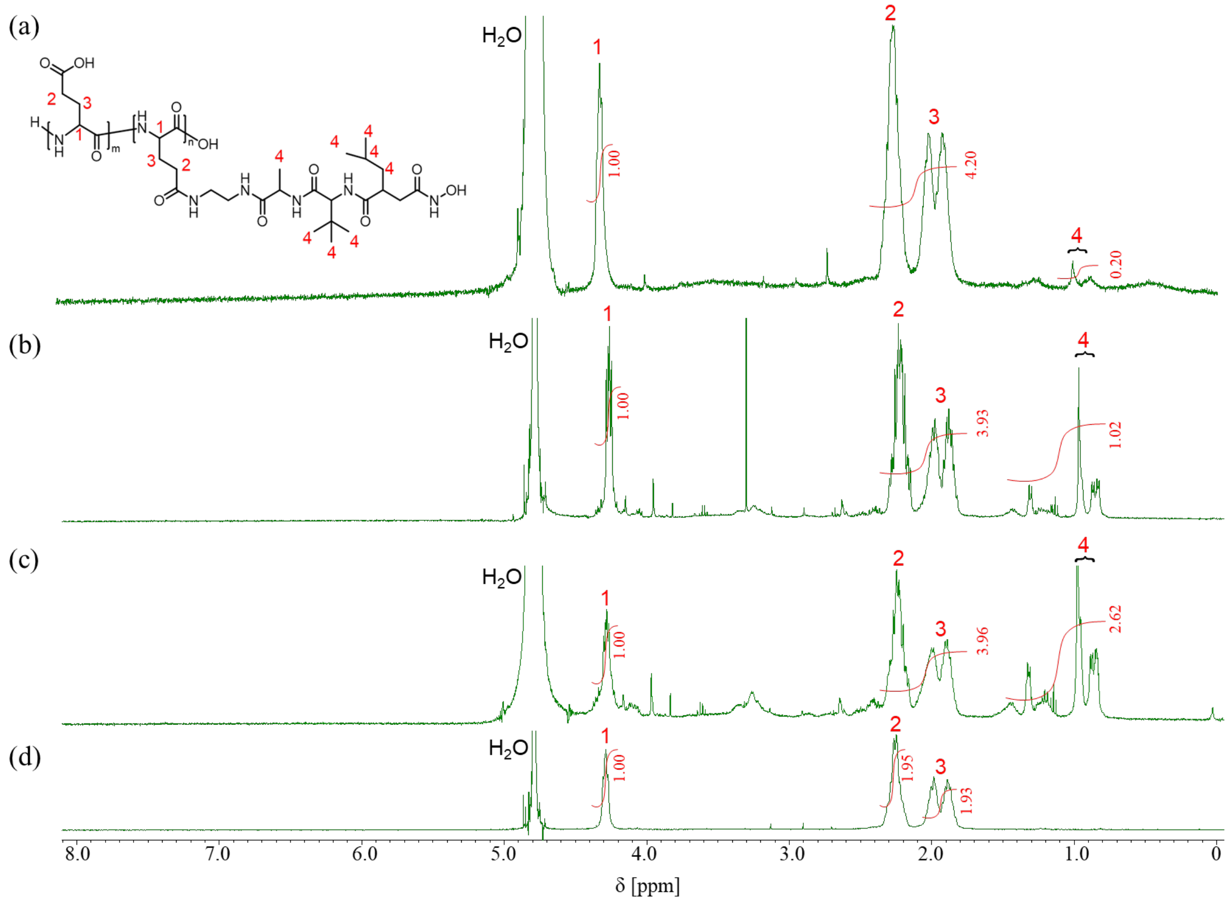


**Figure S3.** [1]H NMR spectra of PGA and PGA-T. The grafting degree of TAPI-2 was calculated from the integration of the peak 1 of PGA and 4 of TAPI-2. (a) 3 mol% (400 MHz, 10 mM NaOD, 25 °C). (b) 10 mol% (400 MHz, 10 mM NaOD, 25 °C). (c) 20 mol% (400 MHz, 10 mM NaOD, 25 °C). (d) PGA (400 MHz, $D_2O$, 25 °C).

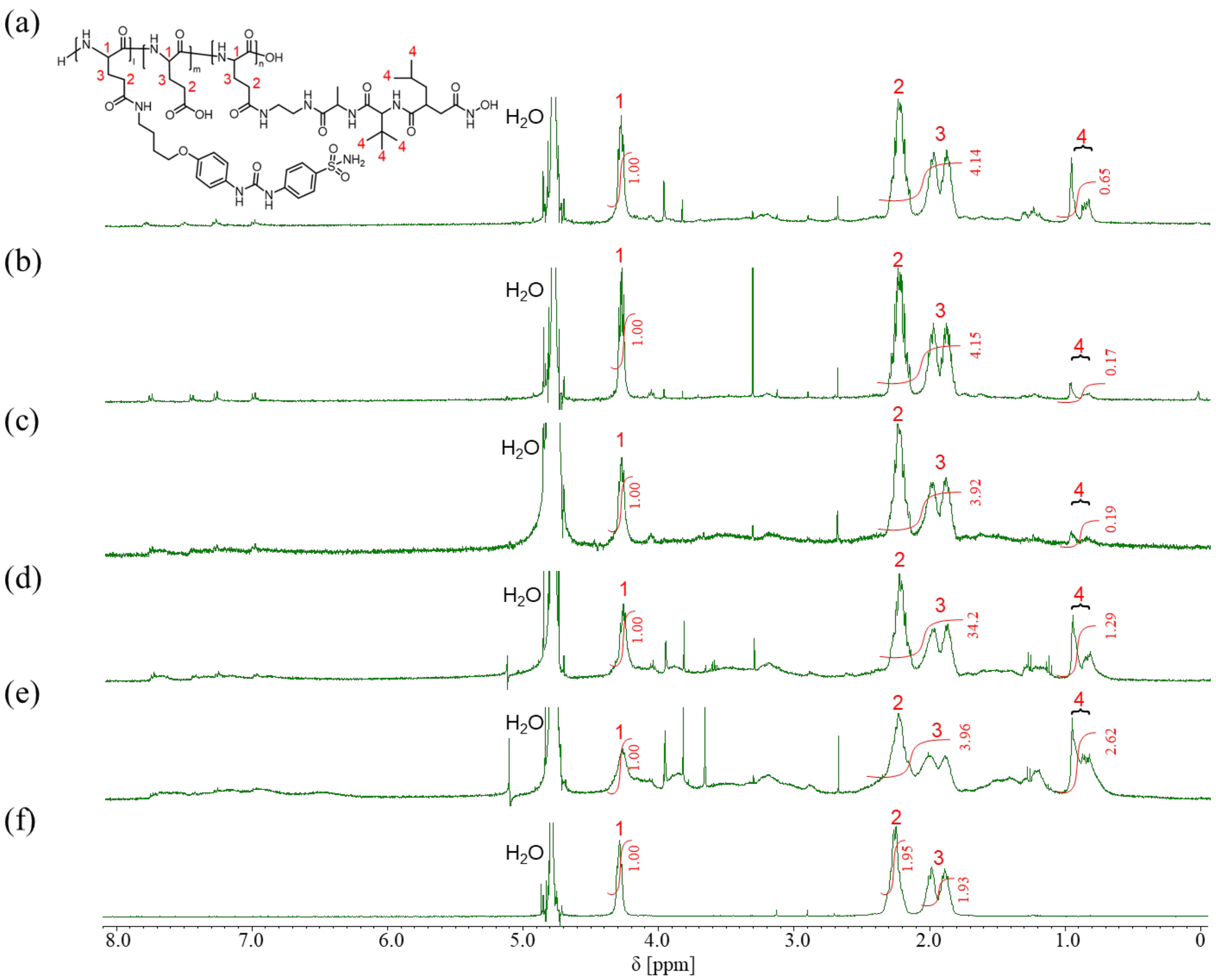


**Figure S4.** $^{1}H$ NMR spectra of PGA and PGA-UT. The grafting degree of TAPI-2 was calculated from the integration of the peak 1 of PGA and 4 of TAPI-2. (a) U-104: 3 mol% and TAPI-2: 10 mol% (400 MHz, 10 mM NaOD, 25 °C). (b) U-104: 3 mol% and TAPI-2: 3 mol% (400 MHz, 10 mM NaOD, 25 °C). (c) U-104: 10 mol% and TAPI-2: 3 mol% (400 MHz, 10 mM NaOD, 25 °C). (d) U-104: 10mol% and TAPI-2: 10 mol% (400 MHz, 10 mM, NaOD, 25 °C). (e) U-104: 20 mol% and TAPI-2: 20 mol% (400 MHz, 10 mM, NaOD, 25 °C). (f) PGA (400 MHz, $D_2O$, 25 °C).

Table S1. The feed ratio for the preparation of PGA derivatives and the grafting degree of U-104, TAPI-2, and ATTO-633 amine.

| | Feed (mol%/ COOH) | | | Feed (eq./ U-104, TAPI-2 or ATTO-633) | Grafting degree (mol%) | | |
|---|---|---|---|---|---|---|---|
| Name | U-104 | TAPI-2 | ATTO-633 | DMT-MM | U-104 | TAPI-2 | ATTO-633 |
| PGA-U19 | 20 | 0 | 0 | 5 | 19 | 0 | 0 |
| PGA-T17 | 0 | 20 | 0 | 5 | 0 | 17 | 0 |
| PGA-U19T15 | 20 | 20 | 0 | 5 | 19 | 15 | 0 |
| PGA-U7 | 10 | 0 | 0 | 5 | 6.6 | 0 | 0 |
| PGA-T6 | 0 | 10 | 0 | 5 | 0 | 6 | 0 |
| PGA-U7T9 | 10 | 10 | 0 | 5 | 7 | 9 | 0 |
| PGA-U7T1 | 10 | 3 | 0 | 5 | 7 | 1 | 0 |
| PGA-U2 | 3 | 0 | 0 | 5 | 2 | 0 | 0 |
| PGA-T1 | 0 | 3 | 0 | 5 | 0 | 1 | 0 |
| PGA-U2T1 | 3 | 3 | 0 | 5 | 2 | 1 | 0 |
| PGA-ATTO | 0 | 0 | 1 | 5 | 0 | 0 | 0.05 |
| PGA-UT-ATTO | 0 | 0 | 1 | 5 | 19 | 15 | 0.01 |

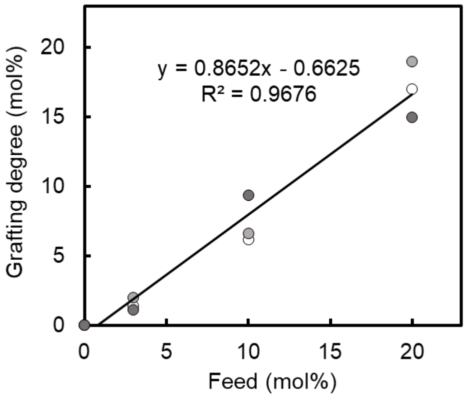


**Figure S5.** Relationship between the feed ratio and the grafting degree of inhibitor. Black: U; white: T; light gray: U of PGA-UT; dark gray: T of PGA-UT. There was a correlation between the feed ratio and the grafting degree.

#### 3-5. Size measurement by Dynamic Light Scattering

The volume averaged hydrodynamic diameter of polymers were measured using dynamic light scattering (in PBS at 37 °C). The concentration of polymers was 1 mg $mL^{-1}$.

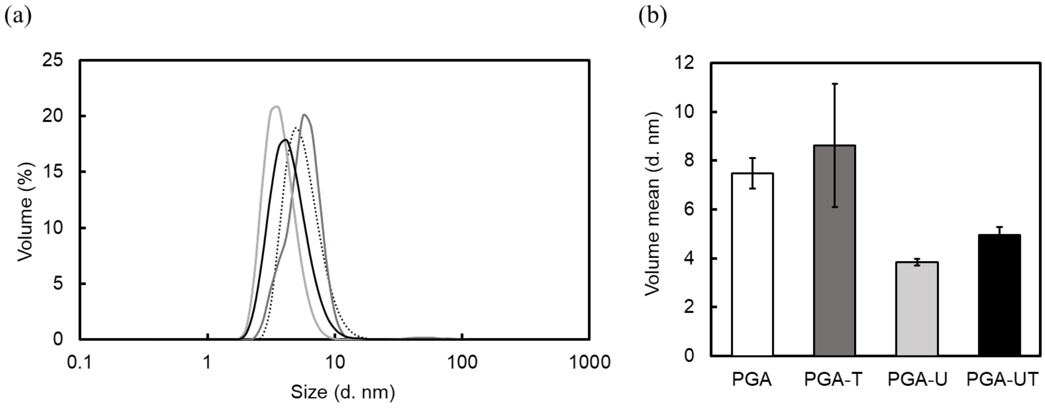


**Figure S6.** (a) Size distribution of synthesized PGA derivatives. Dashed line: PGA; light gray: PGA-U19; dark gray: PGA-T17; and black: PGA-U19T15. (b) Volume means of size of synthesized PGA derivatives.

#### 3-6. Synthesis of fluorescence-labeled PGA derivatives

100 mM carbonate-bicarbonate buffer was prepared by mixing sodium carbonate (0.25 mmol) and sodium hydrogen carbonate in MilliQ (5 mL). PGA (25.0 nmol) was dissolved in 10 mM carbonate-bicarbonate buffer and sonicated 10 min. Then, 500 mM DMT-MM aqueous solution was dropped into the solution and stirred for 10 min. 1 mM ATTO-633 amine MilliQ solution was added into the reaction mixture and stirred for 24 h at room temperature. The reaction mixtures were dialyzed on a dialysis membrane in the following order: against 2.5 mM HCl for 1 day, against 2.5 mM NaOH for 1 day, and against MilliQ for 1 day, respectively, followed by lyophilized to obtain PGA-UT-633. Yield = 100%. The grafting degree of ATTO-633 amine was calculated from the fluorescence intensity at 650 nm.

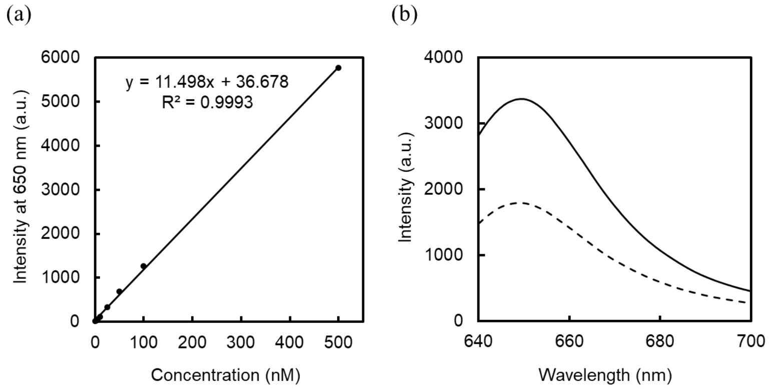


**Figure S7.** Fluorescence measurement in MilliQ. Excitation: 629 nm and Emission: 650 nm. (a) The calibration curve of ATTO-633 amine. (b) Fluorescence spectra of PGA-ATTO (Dashed line: 80 µg $mL^{-1}$) and PGA-UT-ATTO (Gray: 0.4 mg $mL^{-1}$).

### 4. WST assay

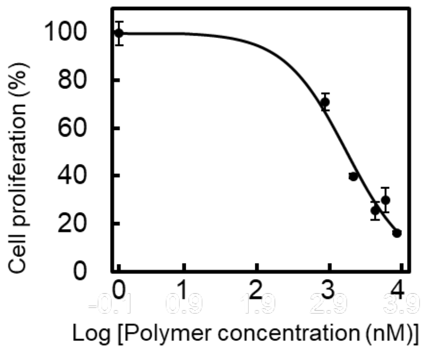


$$Fitting\ equation = \frac{100}{\{1 + 10^{(\log IC_{50} - x)(Hillslope)}\}}$$

**Figure S8.** The $IC_{50}$ fitting curve of PGA-U19T15. The $IC_{50}$ value was 1.33 μM. $R^2 = 0.9842$.

### 5. Visualization of bonding PGA-U19T15 with MDA-MB-231

The obtained images were processed with Imaris Cell Imaging Software 9.0, applied the settings as follows. Algorithm: Enable Region of interest = false, Enable Region Growing = false, Enable Tracking = false. Source Channel: Source Channel Index = 2, Enable Smooth = true, Surface Gain Size = 0.500 μm, Enable Eliminate Background = false, Diameter of Latest Sphere = 4.66 μm. Threshold: Enable Automatic Threshold = false, Manual Threshold Value = 193.576, Active Threshold = true, Enable Automatic Threshold B = true, Manual Threshold Value = 3846.18, Active Threshold B = false. Classify Surfaces: "Number of Voxel Img = 1" above 1.0. We also observed confocal Z slice images (Figure S9). Confocal z slice images suggest that PGA-UT-ATTO might be internalized in the cell but not be in nucleus (Figure S9c, d).[54, 55]

### 6. Competitive Binding Inhibition Assay.

The fluorescence intensity of confocal images was analyzed using the Imaris Cell Imaging Software 9.0 applied the settings as follows. Algorithm: Enable Region of interest = false, Enable Region Growing = false, Enable Tracking = false. Source Channel: Source Channel Index = 2, Enable Smooth = true, Surface Gain Size = 0.500 μm, Enable Eliminate Background = false, Diameter of Latest Sphere = 4.66 μm. Threshold: Enable Automatic Threshold = false, Manual Threshold Value = 190.95, Active Threshold = true, Enable Automatic Threshold B = true, Manual Threshold Value = 3853.46, Active Threshold B = false. Classify Surfaces: "Number of Voxel Img = 1" above 38.

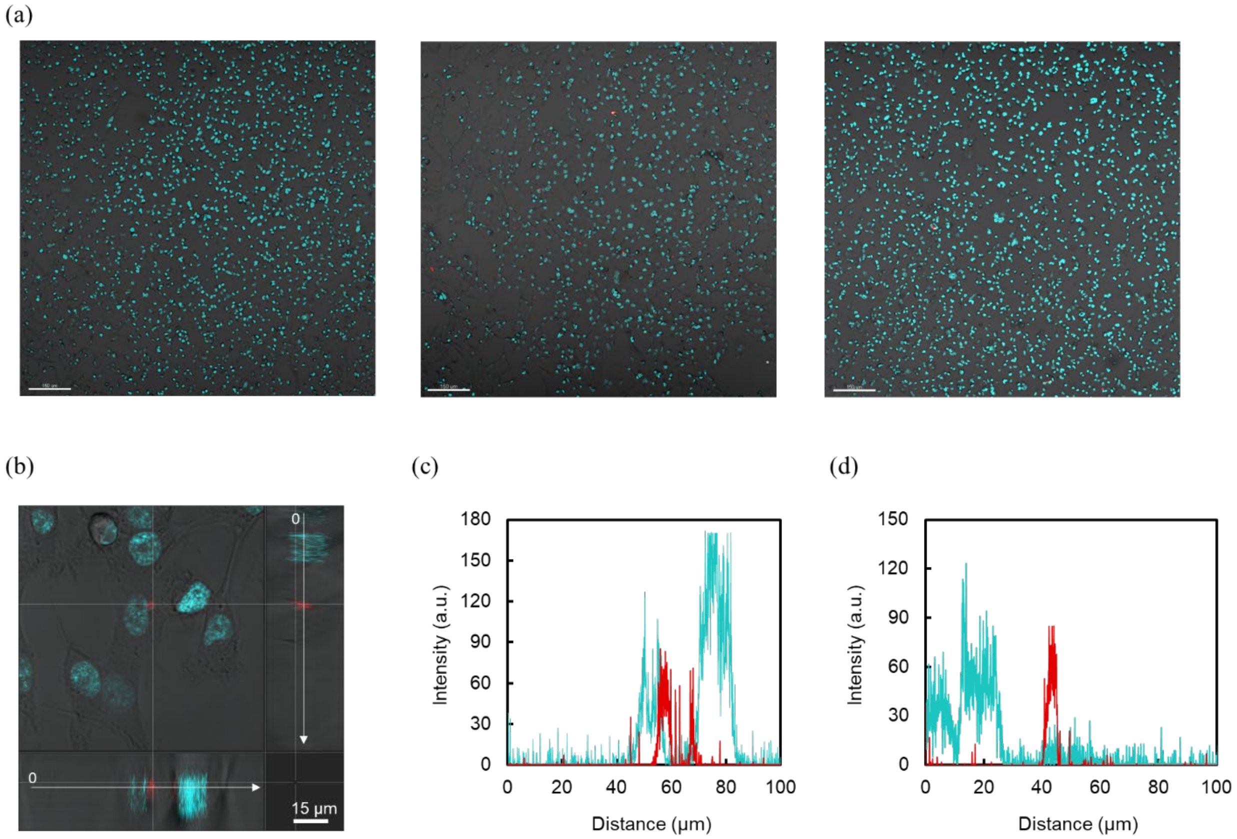


**Figure S9.** (a) Full size confocal images of MDA-MB-231 treated with PGA-ATTO (left), PGA-UT-ATTO (middle), and PGA-UT-ATTO in the presence of U-104 and TAPI-2 for 1 h. Polymer concentration was 100 nM. Red: ATTO-633; blue: Hoechst 33342. Scale bar = 150 µm. (b) Z stack confocal images of MDA-MB-231 treated with PGA-UT-ATTO. Blue: Hoechst 33342; Red: PGA-UT-ATTO. Bottom and right images are xz plane and zy plane, respectively. (c) Plot profiles of the fluorescence in xz plane on the white line in xy plane. (d) Distribution of the fluorescence in zy plane on the white line in xy plane.
